# Meteorin-like (Metrnl) adipomyokine improves glucose tolerance in type 2 diabetes via AMPK pathway

**DOI:** 10.1101/420489

**Authors:** Jung Ok Lee, Hye Jeong Lee, Yong Woo Lee, Jeong Ah Han, Min Ju Kang, Jiyoung Moon, Min-Jeong Shin, Ho Jun Lee, Ji Hyung Chung, Jin-Seok Lee, Chang-Gue Son, Kwon-Ho Song, Tae Woo Kim, Eun-Soo Lee, Hong min Kim, Choon Hee Chung, Kevin R.W. Ngoei, Naomi X.Y. Ling, Jonathan S. Oakhill, Sandra Galic, Lisa Murray-Segal, Bruce E. Kemp, Kyoung Min Kim, Soo Lim, Hyeon Soo Kim

**Author notes:** Corresponding author: Department of Anatomy, Korea University College of Medicine, 73, Inchon-ro, Seongbuk-gu, Seoul 02841, Republic of Korea; E-mail address (H. S. Kim).

## Abstract

Meteorin-like (metrnl) is a recently identified adipomyokine that has beneficial effects on glucose metabolism. However, its underlying mechanism of action is not completely understood. In this study, we have shown that a level of metrnl increase *in vitro* under electrical-pulse-stimulation (EPS) and *in vivo* in exercise mice, suggesting that metrnl is an exercise-induced myokine. In addition, metrnl increases glucose uptake through the calcium-dependent AMPK pathway. Metrnl also increases the phosphorylation of HDAC5, a transcriptional repressor of GLUT4, in an AMPK-dependent manner. Phosphorylated HDAC5 interacts with 14-3-3 proteins and sequesters them in the cytoplasm, resulting in the activation of GLUT4 transcription. The intraperitoneal injection of recombinant metrnl improves glucose tolerance in mice with high fat-induced obesity or type 2 diabetes (db/db), but this is not seen in AMPK β1β2 muscle-specific null mice (AMPK β1β2 MKO). In conclusion, we have demonstrated that metrnl induces beneficial effects on glucose metabolism via AMPK and is a promising therapeutic candidate for glucose-related diseases such as type 2 diabetes.

---

Exercise has the potential to protect against metabolic disease, cardiovascular disease, cancer, and dementia^1^. In response to exercise, skeletal muscle cells secrete various factors called myokines that elicit responses in an auto-, para-, or endocrine manner^2-5^. Myokines are known to improve glucose uptake^6,7^, glucose tolerance^8^, regulation of fat oxidation^8,9^, and satellite cell proliferation^10,11^. Adipose tissue has been established as an endocrine organ that releases various adipokines to control systemic metabolism and energy homeostasis^12,13^. Many contraction-regulated myokines are known to be secreted by adipocytes and are thus called adipomyokines. Adipomyokines are associated with beneficial, exercise-induced metabolic effects^14-16^. We have analyzed adipomyokines such as irisin^17^, fstl-1^18^, resistin^19^, and visfatin^20^. Our findings support the notion that these secreted molecules might mediate some of the well-established, beneficial effects of exercise.

Meteorin-like hormone (metrnl), also known as cometin, subfatin, or IL-39, is a secreted adipomyokine^21,22^. Metrnl is expressed in various tissues, including liver, heart, stromal cells, macrophages, spleen, and central nervous system (CNS) tissues^23,24^. Metrnl expression is increased in white adipose tissue during acute cold exposure, and in muscle tissue after acute bouts of exercise^25^. Metrnl is also known to induce the expression of genes associated with thermogenesis in brown/beige adipocytes, where it stimulates anti-inflammatory cytokines^25^. While the expression and function of metrnl have been explored extensively in fat tissues, the studies on the molecular mechanism of metrnl in skeletal muscle are lacking.

AMP-activated kinase (AMPK) is a master regulator of metabolic homeostasis and an energy-sensing serine/threonine kinase that is activated when cellular energy levels are low^26^. Activation of AMPK is known to stimulate glucose uptake in skeletal muscle, to induce fatty acid oxidation in adipose tissue, and to reduce hepatic glucose production^27^. Activation of AMPK elicits insulin-sensitizing effects during muscle contraction. Skeletal muscle contraction triggers various signaling molecules and AMPK is one of the key proteins in mediating the muscle contraction-induced physiological response. Several myokines (such as IL-15) are regulated by AMPK^28^. We hypothesized that adipomyokines modulate glucose metabolism via muscle contraction-associated molecules, such as AMPK. Here, we tested the hypothesis whether metrnl, a novel exercise induced cytokine improves glucose metabolism in diet induced obese and diabetic mice. Underlying molecular mechanisms responsible for improved glucose homeostasis were also investigated in skeletal muscle cell system and AMPK β1β2-muscle specific null mice.

## Results

### Metrnl levels increased in *in vivo* and *in vitro* muscle contraction models

To address whether the adipomyokine metrnl is affected by muscle contraction, we investigated whether metrnl was upregulated or secreted following muscle contraction. Differentiated myotube L6 cells, an *in vitro* muscle contraction model, underwent electrical pulse stimulation (EPS) to mimic muscle contraction. Metrnl concentrations increased in EPS-conditioned media (Fig 1a). After EPS, metrnl mRNA expression also increased (Fig. 1b). Under EPS conditions, an increased phosphorylation of AMPKα (a key molecule in muscle contraction, Fig. 1c) showed that our experimental system closely mimicked exercise conditions. In our mouse model, metrnl blood concentrations (Fig. 1d) increased after forced treadmill running (1 hr/day for 3 weeks). AMPKα phosphorylation increased in the quadriceps femoris muscles of the exercised mice, showing that exercise activated AMPKα *in vivo* (Fig. 1e). Moreover, glucose tolerance improved after exercise (Fig. 1f, g). Taken together, these results demonstrated that exercise improves the muscle contraction-induced secretion of metrnl.

**Fig 1.**
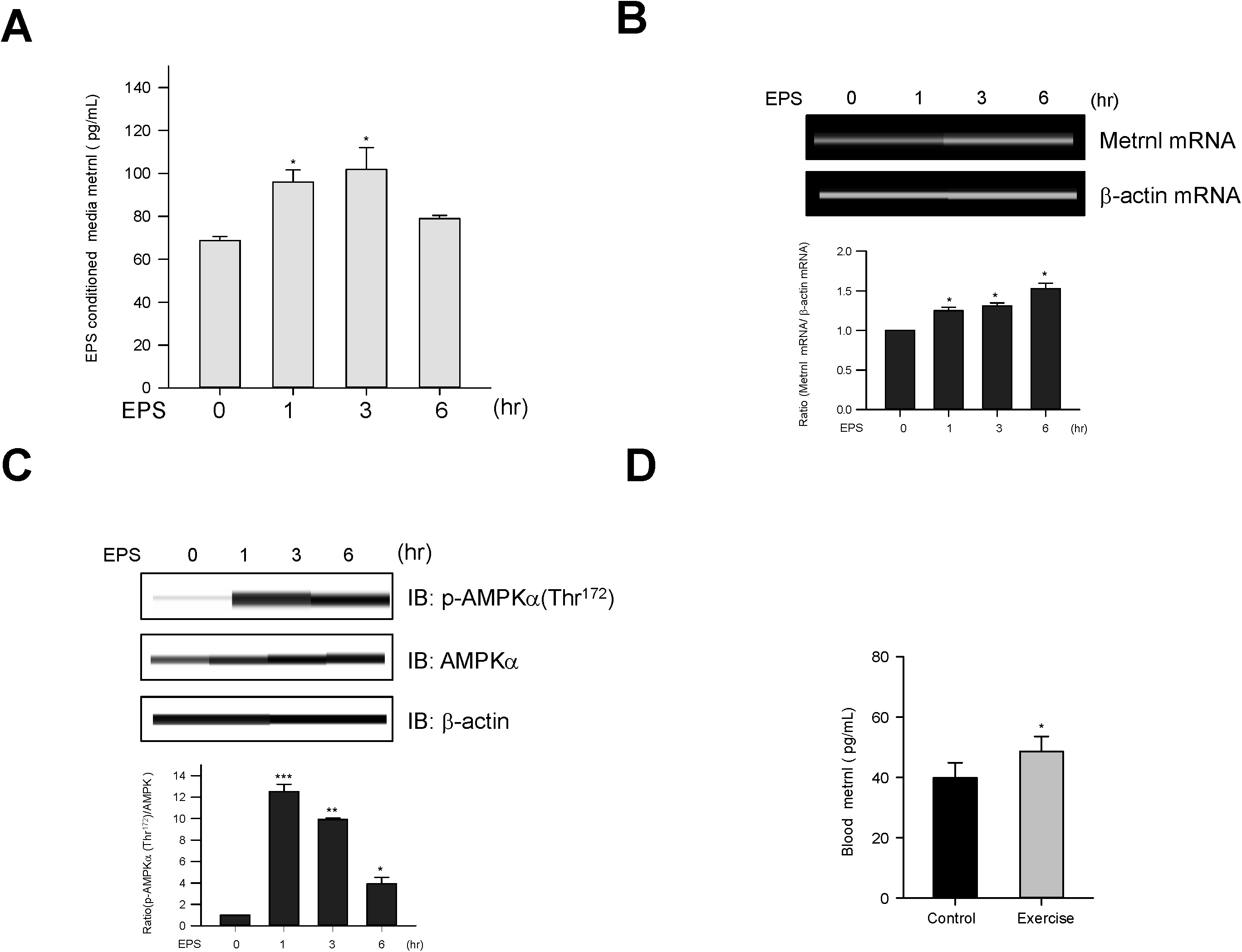
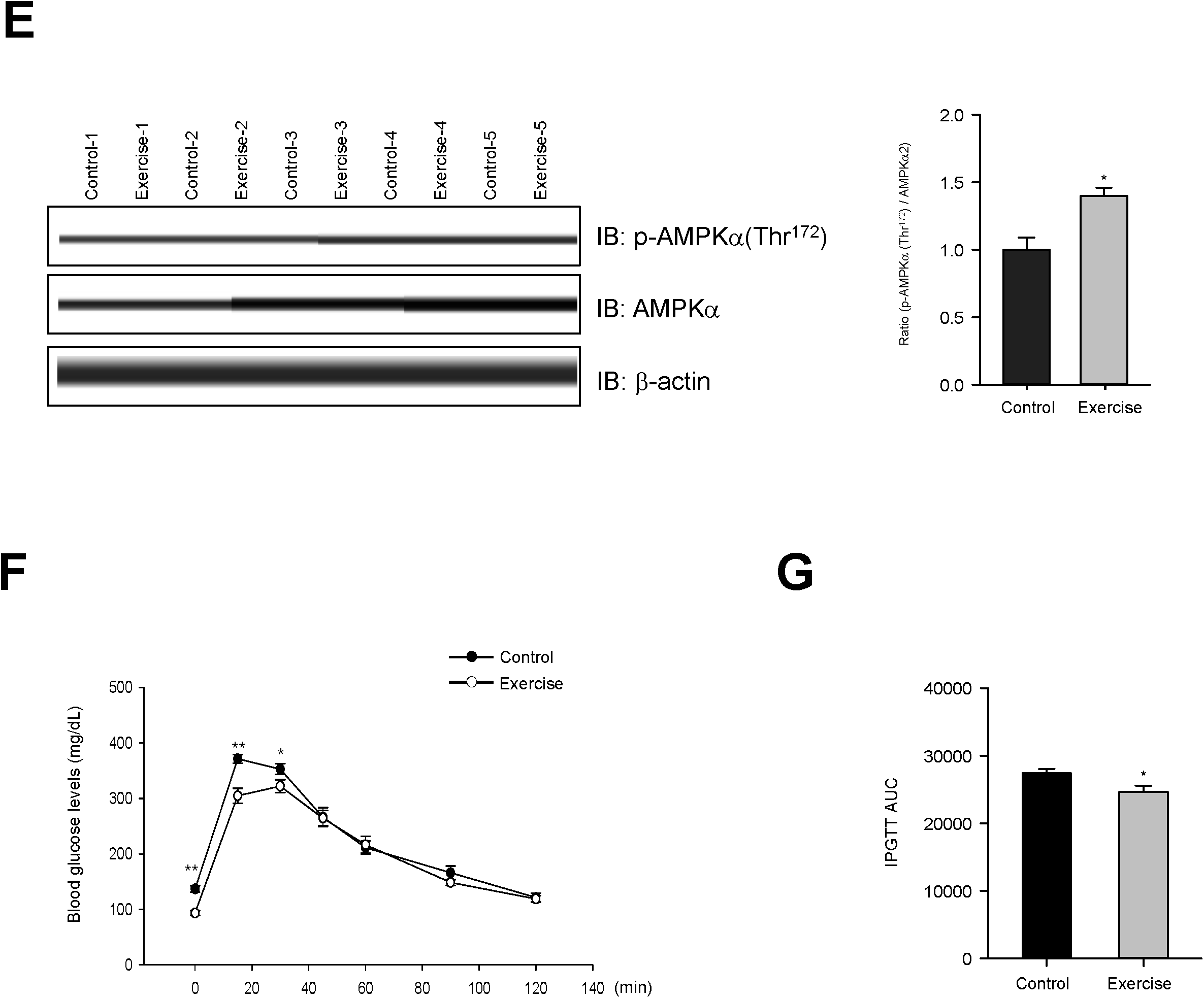
The level of metrnl increased in exercise models *in vivo* and *in vitro*. **a**. L6 myotube cells were subjected to several rounds of EPS (electric pulse stimulation). Lysates were analyzed using a metrnl ELISA kit. **b**. Total mRNA was prepared from L6 myotube cells after electric pulse stimulation, and RT-PCR was performed using metrnl-specific primers. PCR products were separated on a 1% agarose gel and visualized under ultraviolet light, with β-actin as a positive control. **c**. L6 myotube cells were subjected to several rounds of electric pulse stimulation. Lysates were analyzed by Western blotting using an antibody against phospho-AMPKα (Thr^172^); AMPK and β-actin served as a control. **d**. BALB/C mice were divided into groups: sedentary (n=10) and forced treadmill running (n=10). Mice were sacrificed after chronic exercise, and levels of circulating metrnl in blood were measured by ELISA. \**P* < 0.05 compared with control. **e**. Muscle tissues from the same animals were lysed for Western blotting analysis using an antibody against phospho-AMPKα(Thr^172^); AMPKα and β-actin served as controls. F and G. Time course of blood glucose concentrations in sedentary and forced-exercise mice after intraperitoneal injection of glucose (2 mg/kg). Results are displayed as the mean ± SD from three experiments. \**P* < 0.05, \*\**P* < 0.01 and ***P < 0.001 vs. control, as indicated.

### Metrnl stimulates glucose uptake via AMPKα in skeletal muscle cells

To determine whether metrnl affects glucose homeostasis, we evaluated the effect of metrnl on AMPKα phosphorylation in C2C12 mouse skeletal muscle cells. Upon metrnl treatment, AMPKα phosphorylation increased in a dose-dependent (Fig. 2a) and time-dependent (Fig. 2b) manner. Consistent with the increase in AMPKα phosphorylation, the phosphorylation of acetyl-CoA carboxylase (ACC), a downstream substrate of AMPK, also increased after metrnl treatment. Among skeletal muscle cells, differentiated L6 myotubes showed higher glucose uptake than C2C12 cells, suggesting that L6 myotubes were the most promising model for investigating glucose uptake^29^. In the present study, metrnl increased glucose uptake in a dose-dependent manner (Fig. 2c) and up to 3 hr and then reduced (Fig. 2d) in differentiated myotubes. This was suppressed when AMPK was inhibited with compound C or knocked-down with siRNA (Fig. 2e, f). These results suggest that metrnl stimulates glucose uptake via AMPK in skeletal muscle cells.

**Fig 2.**
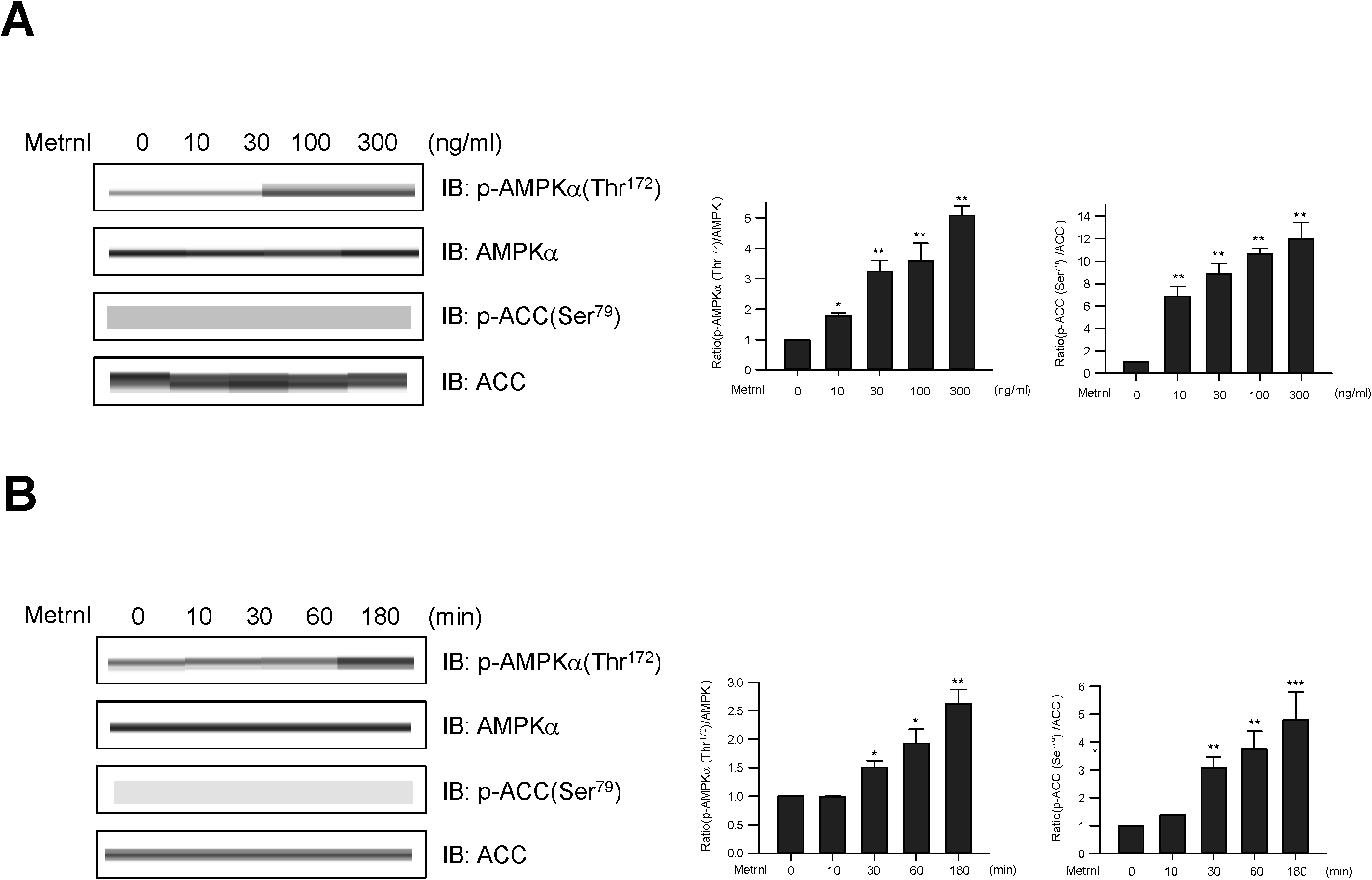
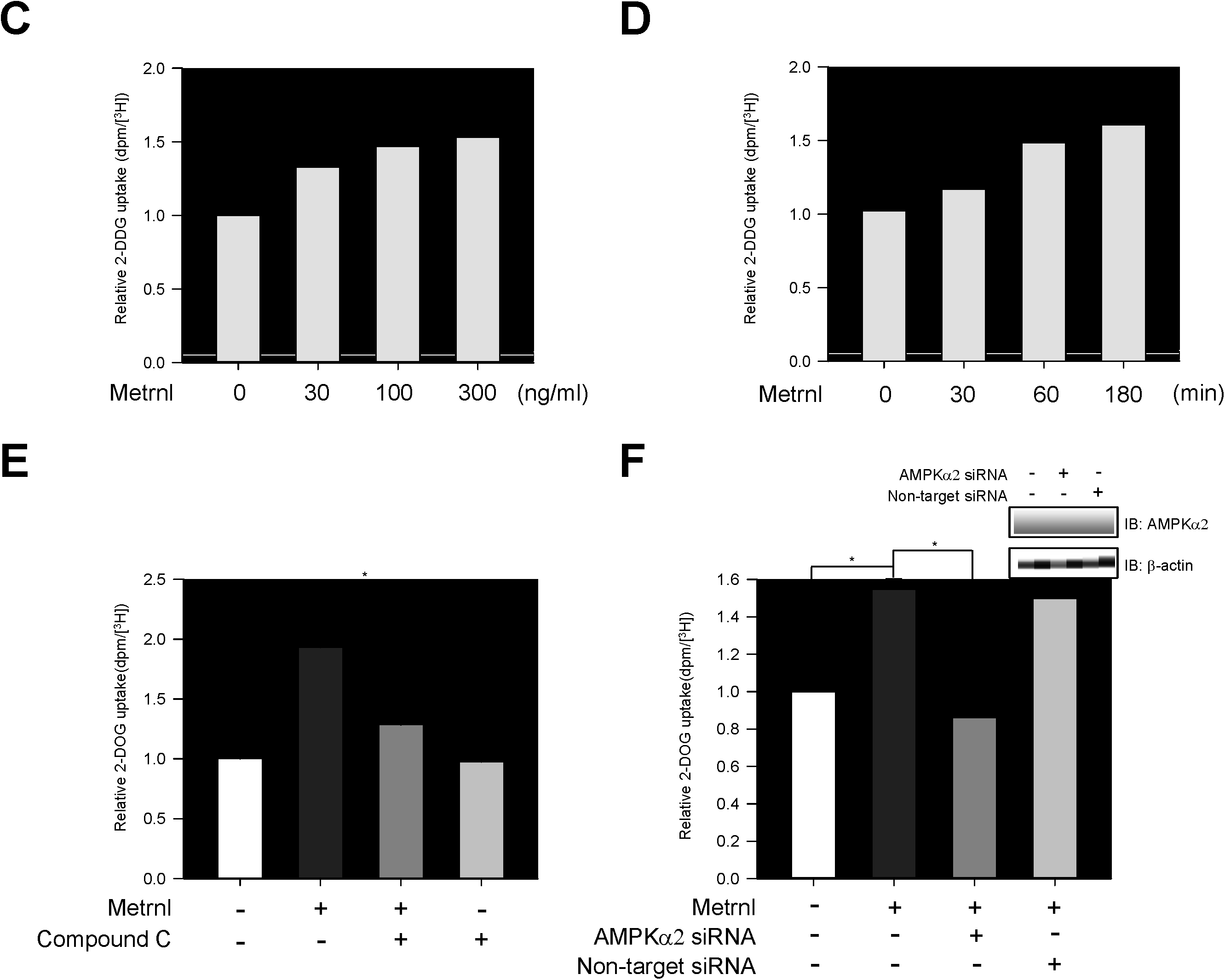
Metrnl stimulates glucose uptake via AMPK in skeletal muscle cells. **a**. Dose-dependent phosphorylation of AMPKα and ACC after metrnl treatment. C2C12 cells were stimulated for 60 min at several concentrations of metrnl. The cell lysates were analyzed by Western blotting using antibodies against phospho-AMPKα(Thr^172^) and phospho-ACC(Ser^79^), while AMPK and ACC served as controls. **b**. Time-dependent phosphorylation of AMPKα and ACC after metrnl treatment. C2C12 cells were incubated with metrnl (100 ng/mL) for the indicated times. Cell lysates were analyzed by Western blotting using antibodies against phospho-AMPKα (Thr^172^) and phospho-ACC (Ser^79^), while AMPK and ACC served as controls. **c**. Dose-dependent uptake of glucose with metrnl treatment. L6 myotube cells were incubated with metrnl at several concentrations for 3 h, and then assayed for glucose uptake. **d**. Time-dependent uptake of glucose with metrnl treatment. L6 myotube cells were incubated with metrnl (100 ng/mL) for the indicated times, and then assayed for glucose uptake. **e**. L6 myotube cells were treated with metrnl (100 ng/mL) for 1 h in the presence of compound C (10 µM), then assayed for glucose uptake. **f**. L6 myotube cells were transiently transfected with AMPKα2 siRNA or non-target siRNA. L6 myotube cells were incubated with metrnl (100 ng/mL) for 1 hour, then assayed for glucose uptake. Results are displayed as the mean ± SD from three experiments. \**P* < 0.05 and \*\**P* < 0.01 vs. control, as indicated.

### Metrnl increases AMPKα phosphorylation by increasing intracellular calcium concentrations

Calcium is a crucial for exercise and contraction induced GLUT4 translocation^30^. We hypothesized that calcium may be involved in metrnl-mediated AMPKα activation. Metrnl increased the fluorescence intensity of cells stained with Fluo-3 AM, a calcium dye (Fig. 3a). Pre-treatment with BAPTA-AM, an intracellular calcium chelator, blocked metrnl-induced AMPKα phosphorylation (Fig 3b). Furthermore, Inhibition using STO-609 of CaMKK2 (calcium/calmodulin-dependent protein kinase 2), a well-known upstream kinase of AMPK, blocked metrnl-induced AMPKα phosphorylation and glucose uptake (Fig. 3c, d). These results demonstrated that metrnl stimulates glucose uptake via calcium-mediated AMPKα phosphorylation.

**Fig 3.**
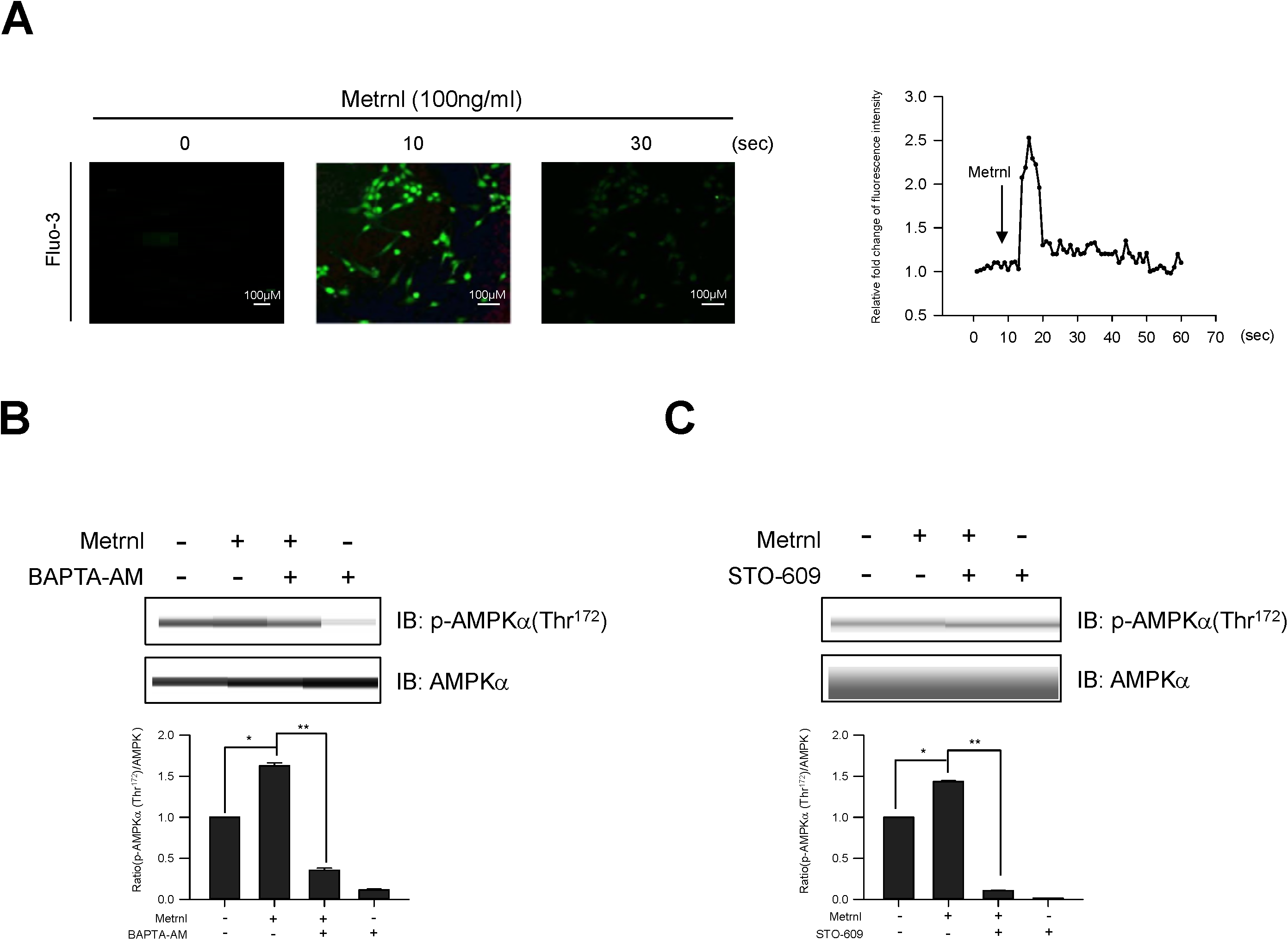
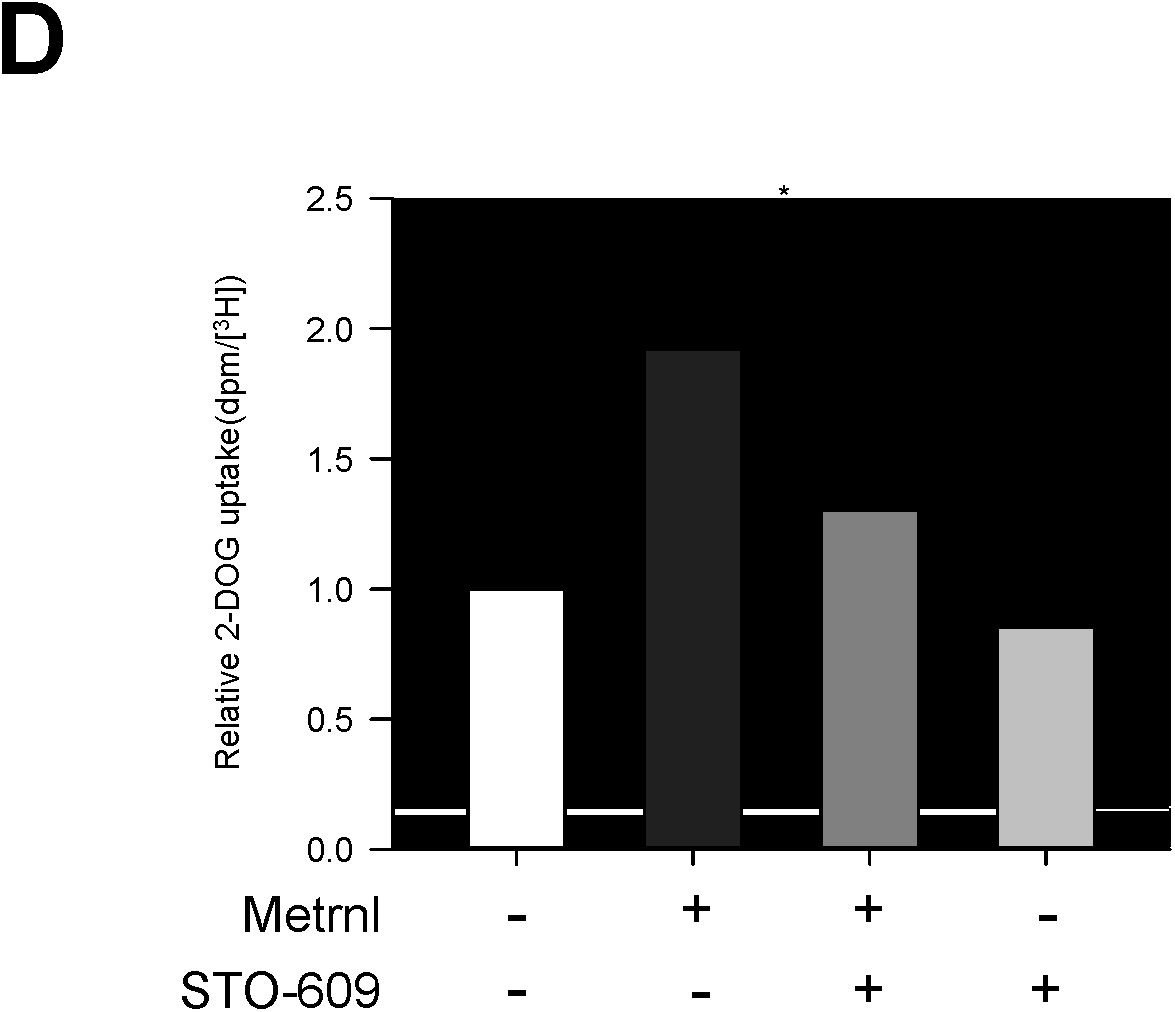
Metrnl activates AMPKα and CaMKK2 by increasing intracellular Ca^2+^. **a**. For Ca^2+^ detection, C2C12 cells were pre-incubated in Fluo-3 AM (10 µM) for 30 min. The Ca^2+^ response was measured after C2C12 cells were pre-treated with calcium-free media and the C2C12 cells were incubated with metrnl (100 ng/mL). The Ca^2+^ concentration correlates with the fluorescence intensity. **b**. C2C12 cells were pre-treated with the membrane-impermeable calcium chelator BAPTA-AM (5 μM), then incubated with metrnl (100 ng/mL) for 60 minutes. Cell lysates were analyzed by Western blotting using an antibody against phospho-AMPKα (Thr^172^), with AMPKα as a control. **c**. C2C12 cells were pre-treated with the CaMKK2 inhibitor STO-609 (5 µM), and then treated with metrnl (100 ng/mL). Cell lysates were analyzed by Western blotting using an antibody against phospho-AMPKα(Thr^172^); AMPKα served as a control. **d**. L6 myotube cells were treated with metrnl (100 ng/mL) for 1 h in the presence of STO-609 (5 µM), then assayed for glucose uptake. \**P* < 0.05 and \*\**P* < 0.01 vs. control, as indicated. Results are displayed as the mean ± SD from three experiments.

### Metrnl increases glucose uptake via the p38 MAPK pathway

Activation of the p38 mitogen-activated protein kinase (MAPK) leads to increased glucose uptake via enhanced GLUT4 translocation^31^. To assess the effect of metrnl on p38 MAPK, we measured the phosphorylation of p38 MAPK after treatment with metrnl. Metrnl increased p38 MAPK phosphorylation in a dose-and time-dependent manner (Fig. 4a, b). We next investigated the role of AMPK in the metrnl-mediated phosphorylation of p38 MAPK. Inhibition or knockdown of AMPK suppressed the metrnl-mediated phosphorylation of p38 MAPK (Fig. 4c, d). To further confirm these findings, we examined the effect of p38 MAPK inhibition on glucose uptake. Inhibition of p38 MAPK with SB202190 blocked metrnl-induced glucose uptake (Fig.4e). These results demonstrated that p38 MAPK is involved in metrnl-mediated glucose uptake, downstream of AMPK.

**Fig 4.**
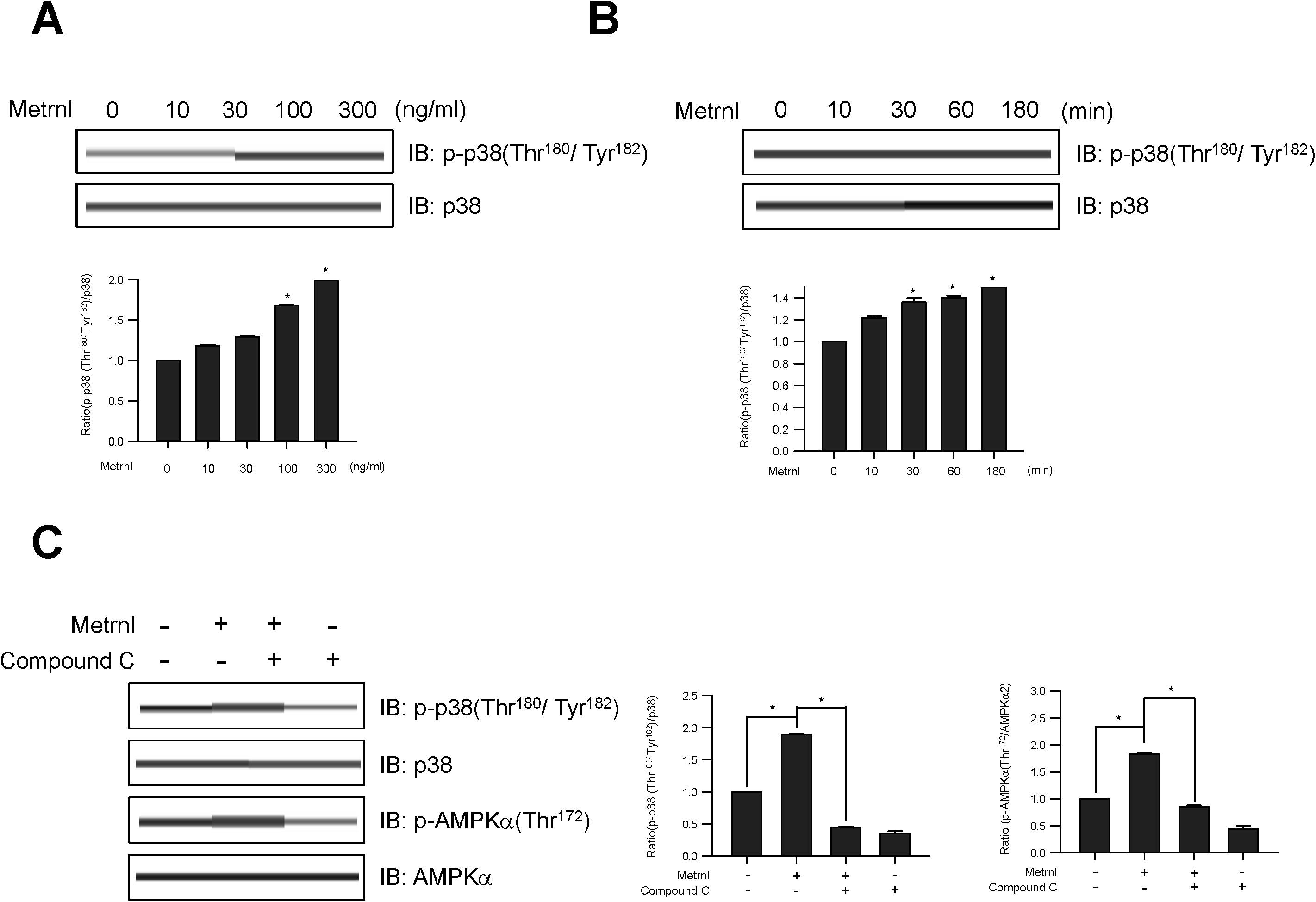
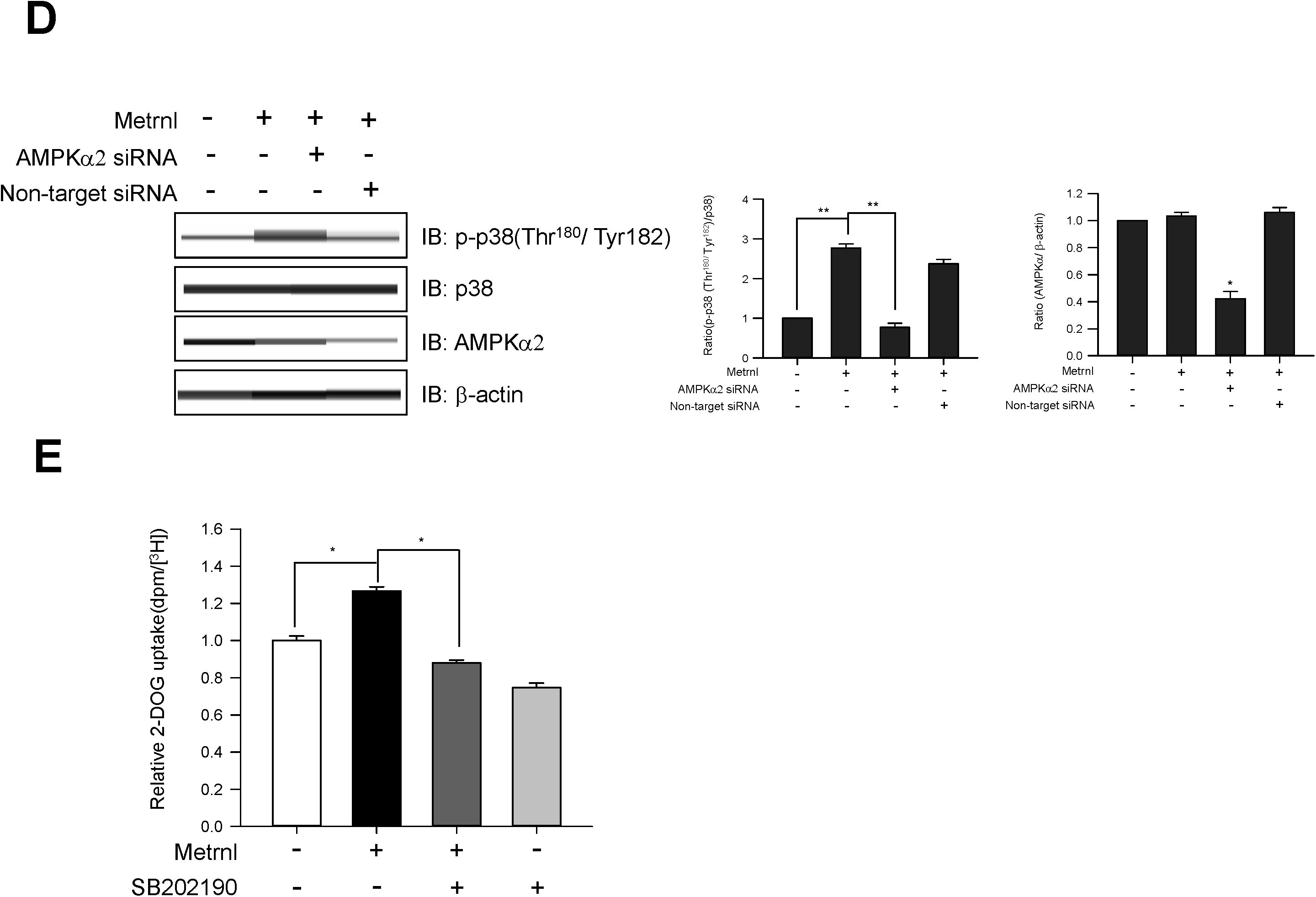
Metrnl increases glucose uptake via the p38 MAPK pathway. **a**. C2C12 cells were stimulated for 60 min with several concentrations of metrnl. The cell lysates were analyzed by Western blotting using antibodies against phospho-p38 MAPK, with p38 MAPK as a control. **b**. Time-dependent phosphorylation of p38 after metrnl treatment. C2C12 cells were incubated with metrnl (100 ng/mL) for the indicated times. Cell lysates were analyzed by Western blotting using antibodies against phospho-p38 MAPK, with p38 MAPK as a control. **c**. C2C12 cells were pre-treated with compound C (30 μM), then treated with metrnl (100 ng/mL). Cell lysates were analyzed by Western blotting using an antibody against phospho-p38 MAPK and phospho-AMPKα (Thr172), with p38 MAPK and AMPKα2 as a control. **d**. C2C12 cells were transiently transfected with AMPKα2 siRNA. Cell lysates were analyzed by Western blotting using an antibody against phospho-p38 MAPK; p38, AMPKα and β-actin served as a control. **e**. L6 myotube cells were treated with metrnl (100 ng/mL) for 1 h in the presence of SB202190 (20 µM), then assayed for glucose uptake. Results are displayed as the mean ± SD from three experiments. \**P* < 0.05 and \*\**P* < 0.01 vs. control, as indicated.

### Metrnl increases GLUT4 expression by stimulating the phosphorylation of HDAC5

Glucose transporter 4 (GLUT4) is essential for glucose uptake in peripheral tissues^32^ and is known to be largely regulated at the transcriptional level^33^. Histone deacetylase 5 (HDAC5) is a corepressor of GLUT4 transcription, being exported from the nucleus after phosphorylation ^33^, while 14-3-3 chaperone protein is known to mediate the nuclear export of HDAC5^35^. In the current study, we showed that metrnl increased the expression of GLUT4 mRNA (Fig. 5a) and GLUT4 protein (Fig. 5b). In addition, metrnl increased HDAC5 phosphorylation (Fig. 5c). This increase in phosphorylation was not observed upon inhibition or knockdown of AMPKα2 (Fig. 5d and e), suggesting that metrnl phosphorylates HDAC5 through AMPKα. We also confirmed that metrnl induced the cytosolic translocationof phosphorylated HDAC5 by cytosolic fractionation and immunocytochemistry (ICC) experiments (Fig. 5f and g). Metrnl also increased the interaction between phosphorylated HDAC5 and 14-3-3 in both immunoprecipitation (IP) and ICC experiments (Fig. 5h and 5i), suggesting that 14-3-3 helps to sequester HDAC5 in the cytoplasm. To confirm whether metrnl affects HDAC5 binding to the GLUT4 promoter, chromatin immunoprecipitation (ChIP) assays were performed. Notably, metrnl treatment reduced HDAC5 binding to the GLUT4 promoter region. This was accompanied by an increase in histone H3 acetylation at the same region (Fig. 5j). These data showed that metrnl upregulates GLUT4 expression by releasing HDAC5 from its promoter region.

**Fig 5.**
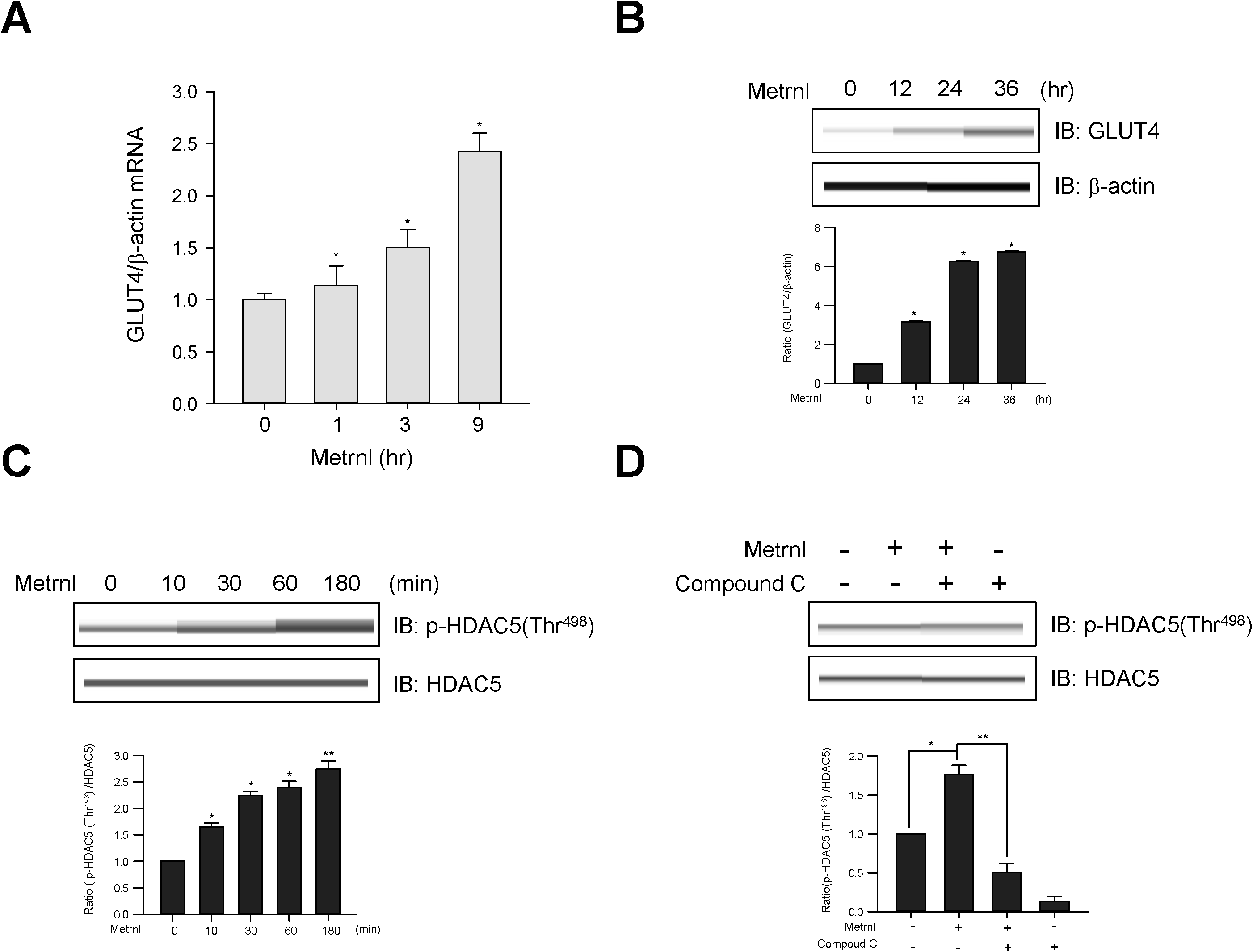
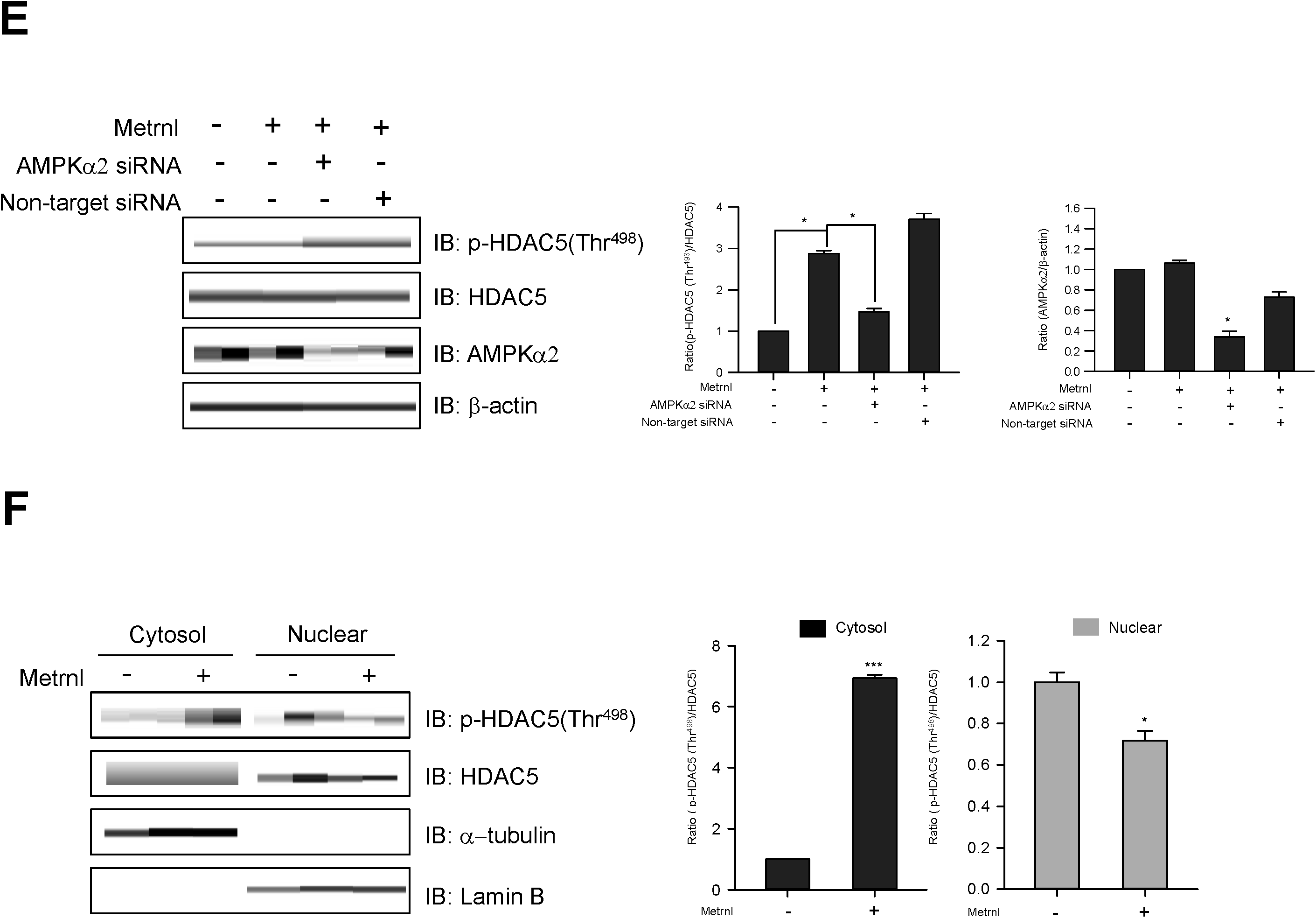
Metrnl increases GLUT4 expression by stimulating HDAC5 phosphorylation. **a**. Total mRNA from C2C12 cells were prepared after metrnl (100 ng/mL) treatment for the indicated times, and real-time qRT-PCR was performed using GLUT4-specific primers. For these assays, β-actin mRNA served as a positive control. **b**. C2C12 cells were stimulated with metrnl (100 ng/mL) for the indicate4d times. The cell lysates were analyzed by Western blotting using antibodies against GLUT4, with β-actin as a control. **c**. Time-dependent phosphorylation of HDAC5 after metrnl treatment. C2C12 cells were incubated with metrnl (100 ng/mL) for the indicated times. Cell lysates were analyzed by Western blotting usingantibodies against phospho-HDAC5 (Thr^498^), while HDAC5 served as controls. **d**. C2C12 cells were pre-treated with compound C (30 μM), then treated with metrnl (100 ng/mL). Cell lysates were analyzed by Western blotting using an antibody against phospho-HDAC5 (Thr^498^); HDAC5 served as a control. **e**. C2C12 cells were transiently transfected with AMPKα2 siRNA or non-target siRNA. Cell lysates were analyzed by Western blotting using antibodies against phospho-HDAC5 (Thr^498^) and AMPKα2; HDAC5 and β-actin served as controls. **f**. C2C12 cells were treated with metrnl (100 ng/mL). Cytosolic and nuclear proteins were extracted from the cells. The phosphorylation of HDAC5 was evaluated by Western blot analysis. HDAC5 served as a control. Western blotting was performed on nuclear and cytosolic fractions to detect nuclear (lamin B) and cytosolic (α-tubulin) marker proteins. **g**. Representative images of phospho-HDAC5 treated with metrnl for 30?min. Scale bars, 10?μm. **h**. C2C12 cells were immunoprecipitated with anti-14-3-3 antibody, followed by Western blotting using anti-phospho-HDAC5. **i**. Representative images (phospho-HDAC5 and 14-3-3 objective images) of cells treated with metrnl for 1?h. Scale bars, 10?μm. **j**. The relative occupancy of HDAC5 and AcH3 on the GLUT4 promoter was assessed by qChIP analyses following 60 min of metrnl (100 ng/mL) treatment. The ChIP data represent IP values for each region’s ratio relative to the input. The results shown are from three independent experiments. \**P* < 0.05 and \*\**P* < 0.01 vs. control, as indicated. Results are displayed as the mean ± SD from three experiments.

### Metrnl stimulates the translocation of GLUT4 by AMPKα-induced TBC1D1 phosphorylation

TBC1D1 is a Rab-GTPase-activating protein (GAP) known to be phosphorylated in response to AMPK activation and involved in GLUT4 trafficking^36,37^. We therefore tested the possible involvement of TBC1D1 in metrnl-mediated glucose regulation. Metrnl induced TBC1D1 (Ser^237^) phosphorylation in a time-dependent (Fig. 6a) and dose-dependent (Fig. 6b) manner. An increase in TBC1D1 phosphorylation was not observed when AMPKα was inhibited or knocked down (Fig. 6c and d). To provide further evidence of this, we performed membrane fractionation and ICC experiments. Metrnl increased GLUT4 translocation to the plasma membrane in both experiments (Fig. 6e and f). Insulin was used as positive control for GLUT4 translocation. A colorimetric assay was used to measure cell surface localization of GLUT4myc. Metrnl increased plasma membrane GLUT4myc in time dependently (Fig. 6G). The inhibition and down-regulation of AMPKα2 was not observed plasma membrane GLUT4myc in the presence of metrnl (Fig. 6H and I). However, the down-regulation of TBC1D1 increased the translocation of GLUT4myc into the plasma membrane (Fig. 6J). These results suggest that metrnl stimulates GLUT4 translocation through the AMPKα-TBC1D1 signaling pathway.

**Fig 6.**
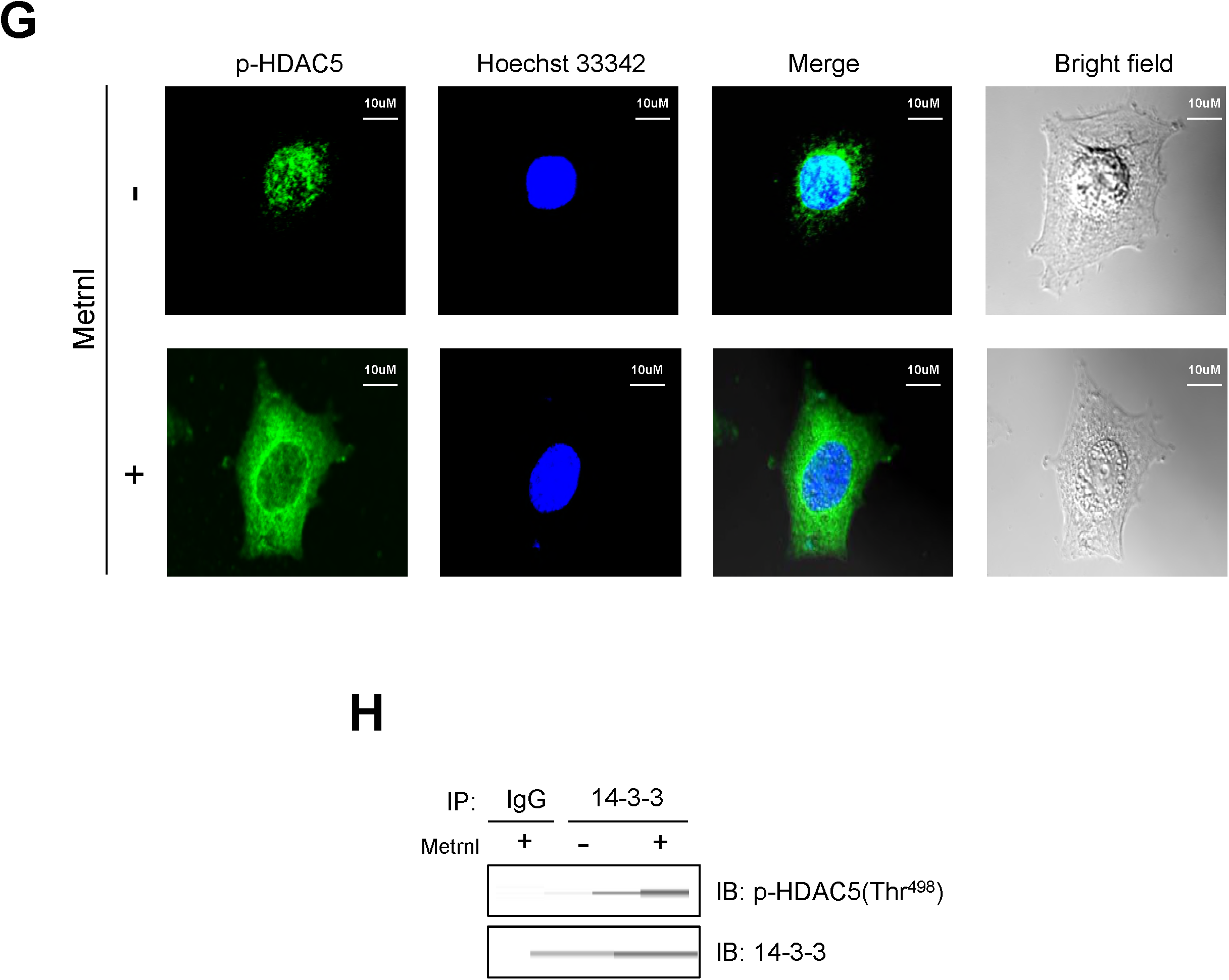
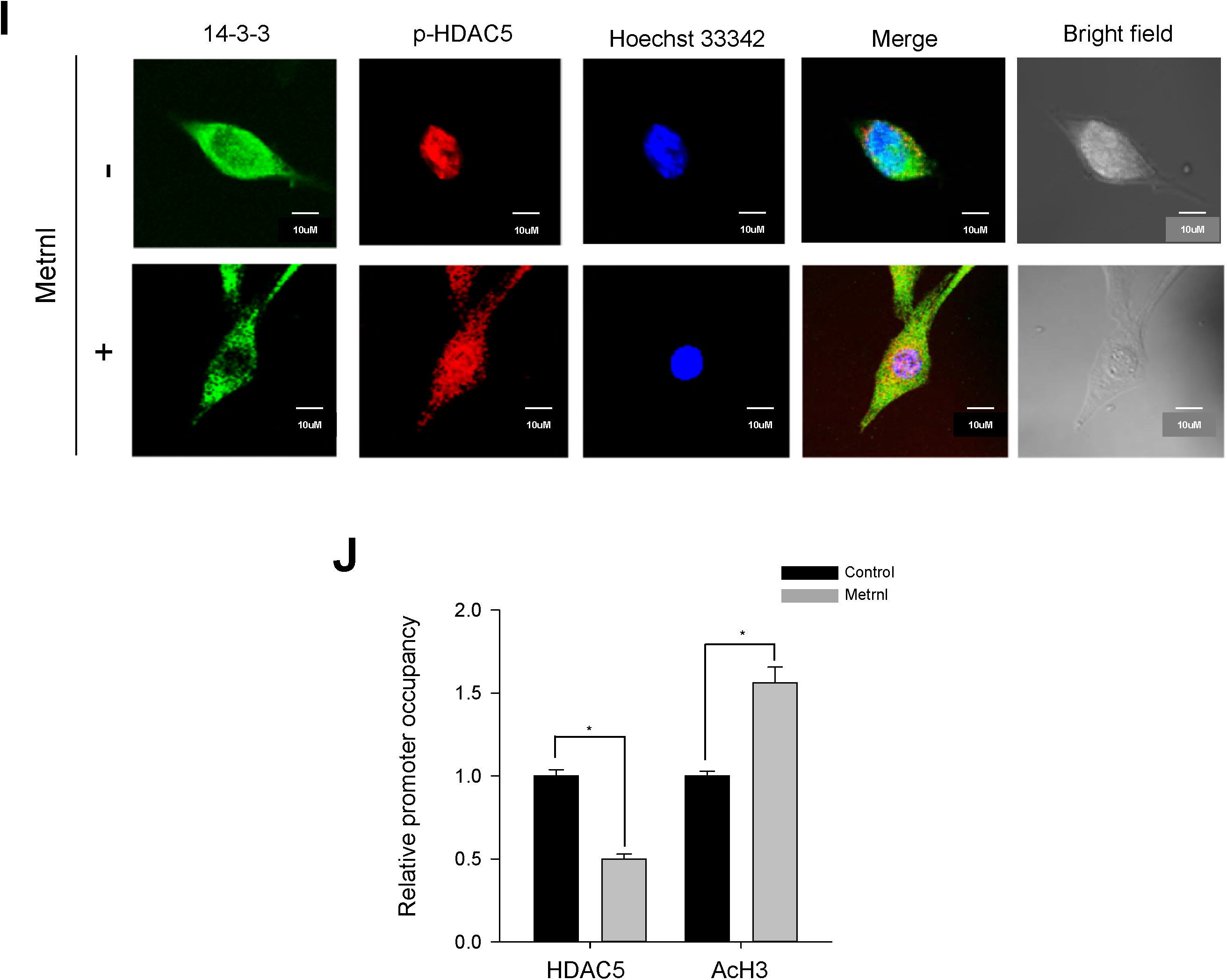
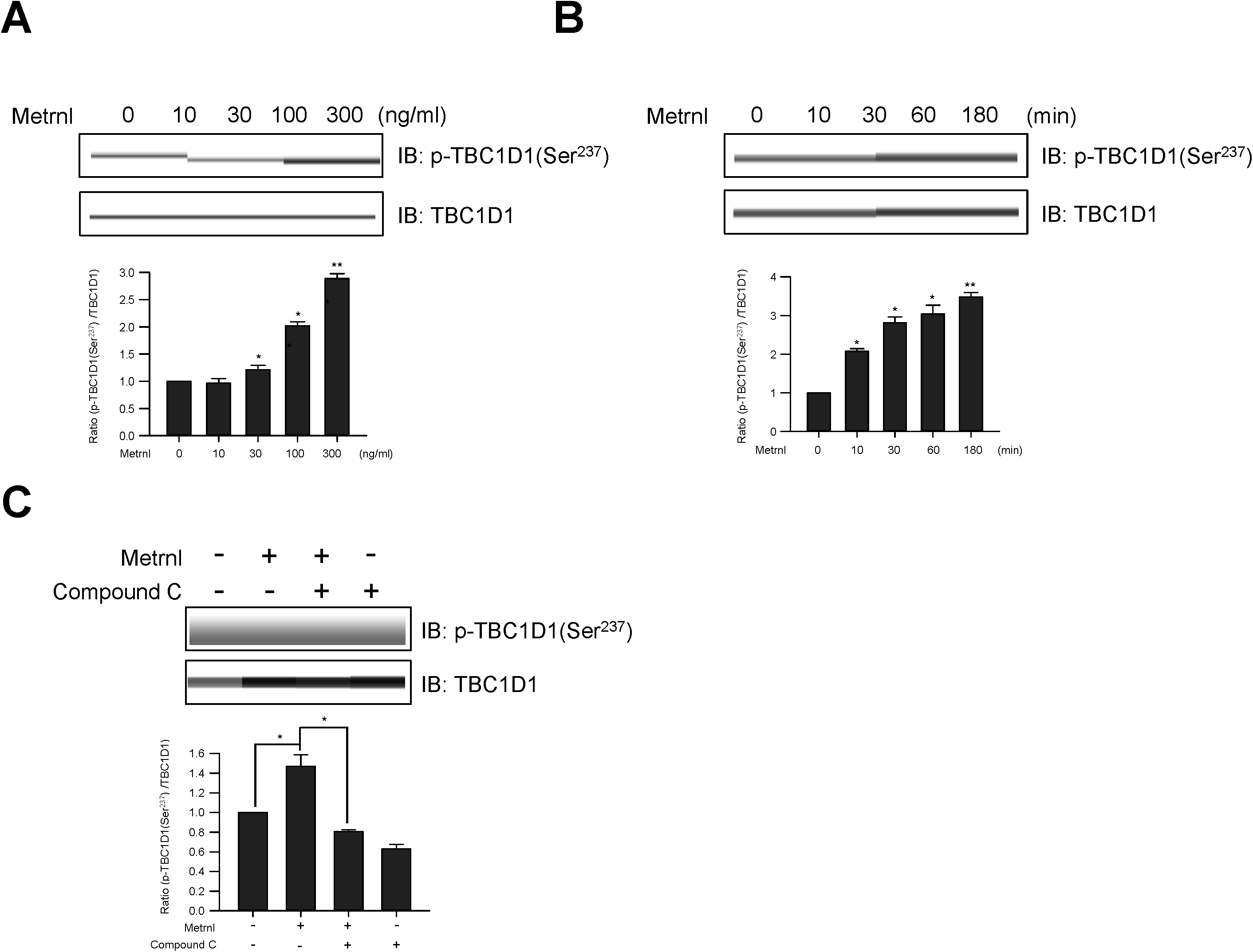
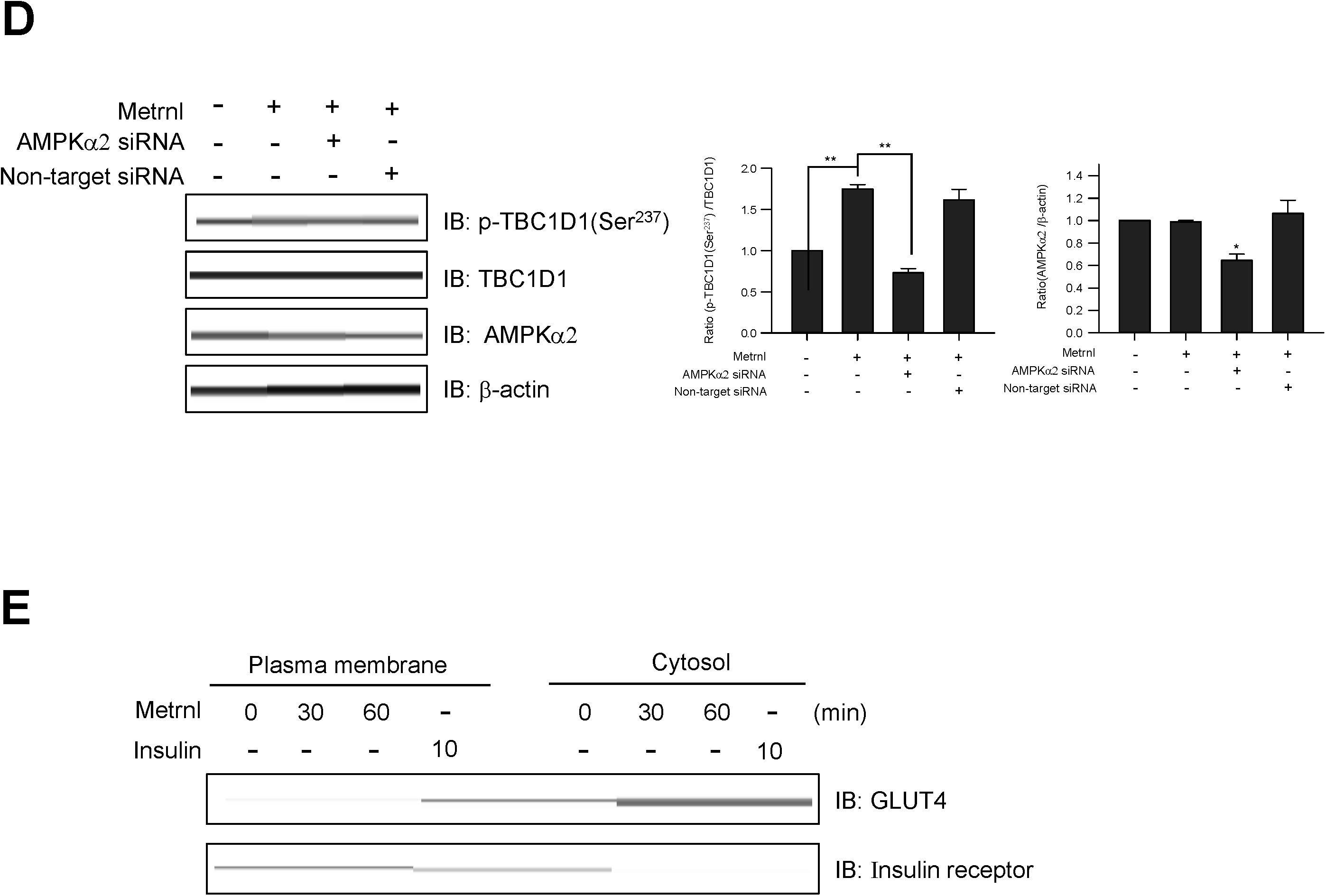
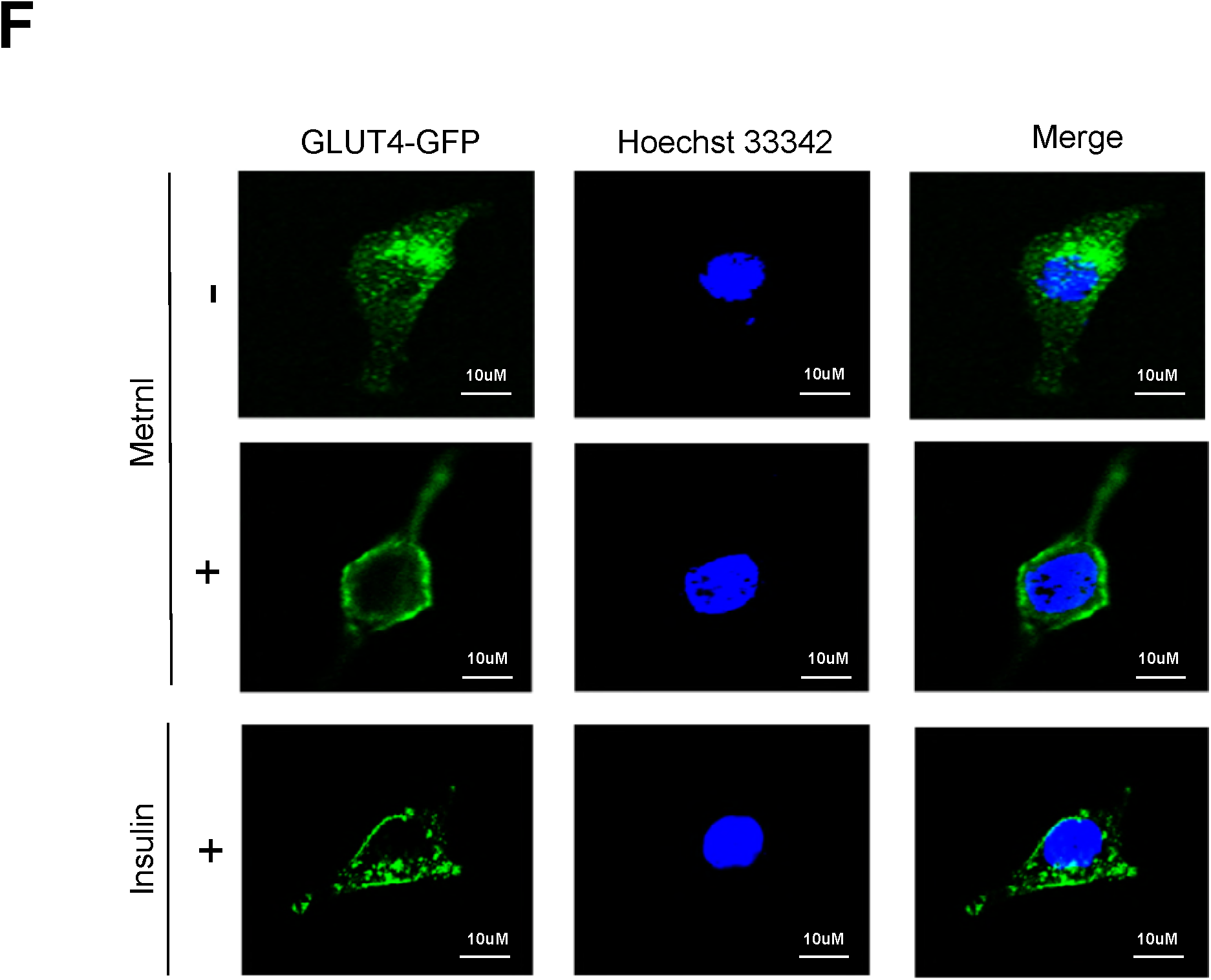
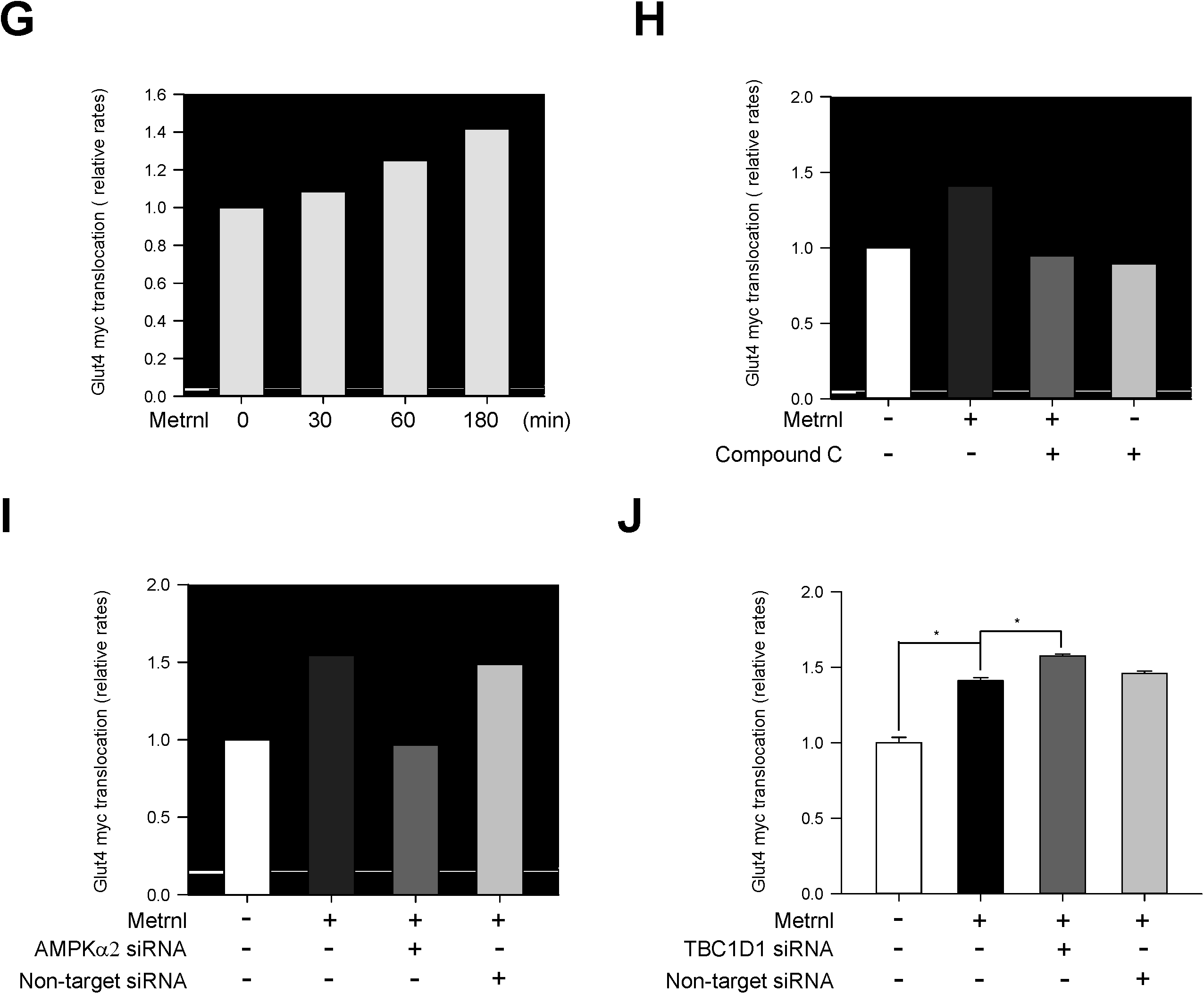
Metrnl stimulates GLUT4 translocation by AMPK-induced TBC1D1 phosphorylation. **a**. C2C12 cells were stimulated for 1 h with different concentrations of metrnl. **b**. C2C12 cells were incubated with metrnl (100 ng/mL) for the indicated times. Cell lysates were analyzed by Western blotting using an antibody against phospho-TBC1D1 (Ser^237^); TBC1D1 served as a control. **c**. C2C12 cells were pre-treated with compound C (10 μM), then treatedwith metrnl (100 ng/mL). Cell lysates were analyzed by Western blotting using an antibody against phospho-TBC1D1 (Ser^237^); TBC1D1 served as a control. **d**. C2C12 cells were transiently transfected with AMPKα2 siRNA or non-target siRNA. Cell lysates were analyzed by Western blotting using antibodies against phospho-TBC1D1 (Ser^237^) and AMPKα2; TBC1D1 and β-actin served as controls. **e**. Fractionated plasma membranes from C2C12 cells were pre-treated with metrnl (100 ng/mL) or insulin (100 nM). Plasma membrane (PM) proteins were analyzed by Western blotting using an antibody against GLUT4; insulin receptor (IR) served as a plasma membrane marker, and β-actin served as a control. **f**. Representative images (GLUT4, Hoechst, and merged) of cells treated with metrnl for 1 h. Insulin (100 nM) was used as a positive control. Scale bars, 10 μm. **g**. Dose-dependent expression of GLUT4myc with metrnl treatment. L6 myotube cells were incubated with metrnl at several concentrations for 3 h, and then cell surface expression of GLUT4myc was detected using an antibody-coupled colorimetric absorbance assay. **h**. L6 myotube cells were treated with metrnl (100 ng/mL) for 1 h in the presence of Compound C (10 µM), then cell surface expression of GLUT4myc was detected using an antibody-coupled colorimetric absorbance assay. **i and j.** L6 myotube cells were transiently transfected with both AMPKα2 and TBC1D1 siRNA for 48 h prior to metrnl (100 ng/mL) for 1 hr. cell surface expression of GLUT4myc was detected using an antibody-coupled colorimetric absorbance assay. Results are displayed as the mean ± SD from three experiments. \**P* < 0.05 and \*\**P* < 0.01 vs. control, as indicated.

### Metrnl stimulates AMPKα phosphorylation and glucose uptake in mouse primary skeletal muscle cells

To provide physiological relevance, we next investigated the effects of metrnl in prepared primary myoblast cells. Primary myoblasts were prepared from quadriceps femoris tissues of wild-type mice (BALB/c). Metrnl significantly increased calcium levels in primary myoblasts, with maximum fluorescence detected at 10 secs (Fig.7a). In addition, metrnl increased the phosphorylation of AMPKα and its downstream target ACC in a time-dependent manner (Fig. 7b). In addition, metrnl increased glucose uptake in differentiated primary myotubes also (Fig. 7c), supporting the biologically relevance of metrnl.

**Fig 7.**
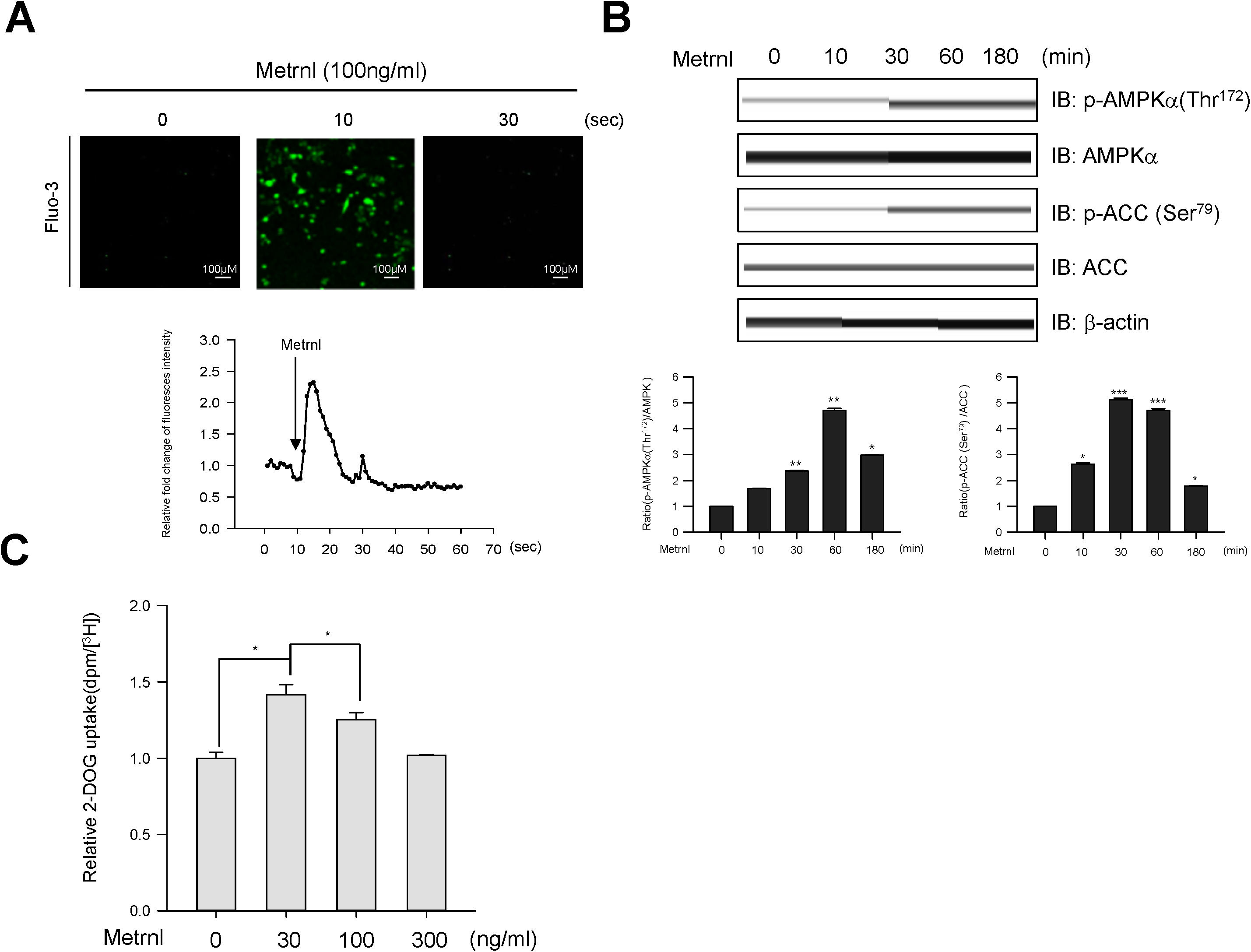
Metrnl regulates AMPK phosphorylation and glucose uptake in mouse primary skeletal muscle cells. **a**. For Ca^2+^ detection, myoblasts were pre-incubated with Fluo-3 AM (5 μM) for 30 minutes. The Ca^2+^ response was measured after treatment with metrnl using a confocal microscope. Scale bars, 100 μm. **b**. Mouse primary skeletal muscle cells were stimulated withmetrnl for the indicated times. Cell lysates were analyzed by Western blotting using antibodies against phospho-AMPKα (Thr^172^) and phospho-ACC (Ser^79^); AMPKα and ACC served as controls. **c**. Dose-dependent uptake of glucose with metrnl treatment. Primary myotube cells were incubated with metrnl at several concentrations for 1 h, and then assayed for glucose uptake. Results are displayed as the mean ± SD from three experiments. \**P* < 0.05 and \*\**P* < 0.01 vs. control, as indicated.

### The administration of metrnl improved glucose tolerance in mice

To investigate the effect of metrnl on glucose tolerance *in vivo*, we prepared both recombinant GST-tagged metrnl protein and GST protein purified from *E. coli* (Fig. 8a). Upon GST –metrnl treatment, AMPKα phosphorylation increased in C2C12 cells (Fig. 8b), confirming the biological activity of the recombinant protein. To examine the effect of metrnl *in vivo*, we administered recombinant metrnl to C57BL/6 mice (n=12 per group). Intraperitoneal injection of recombinant metrnl downregulated blood glucose levels and improved glucose tolerance (Fig. 8c and d). To confirm the effect of metrnl in a disease model, we used type 2 diabetic (db/db) mice. Administration of metrnl ameliorated the impaired glucose tolerance in db/db mice (Fig. 8e and f) and lowered blood glucose levels (Fig. 8g). The phosphorylation of AMPKα increased in the extensor digitorum longus (EDL) tissue of metrnl-treated mice (Fig. 8h), suggesting that metrnl improves glucose tolerance in db/db mice by activating AMPKα in skeletal muscle tissue. Regarding chronic effect of metrnl on glucose tolerance in diet induced obese mice, intraperitoneal injection of recombinant metrnl significantly improved glucose tolerance in both normal chow diet and high fat diet group (Fig. 8i and j).

**Fig 8.**
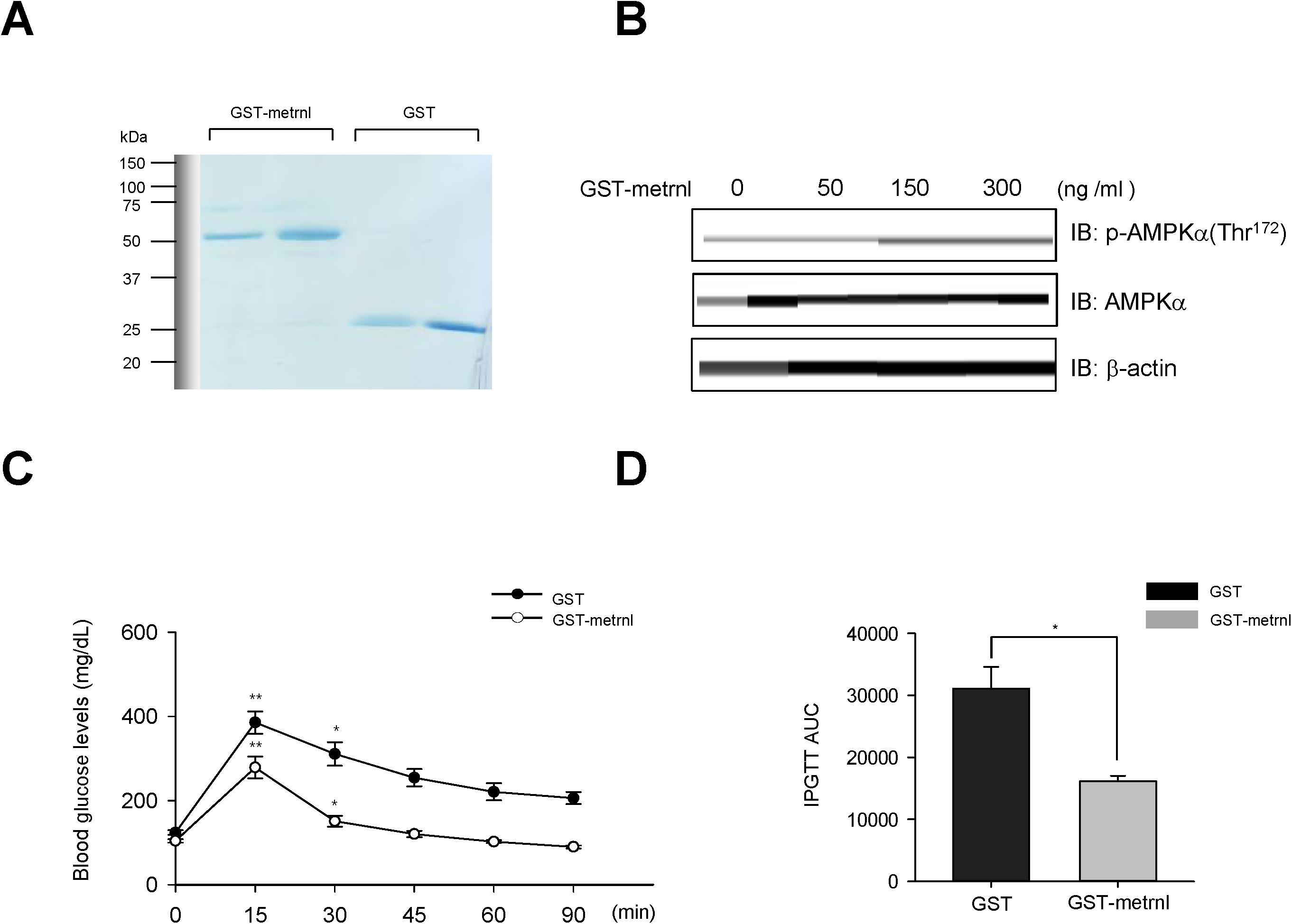
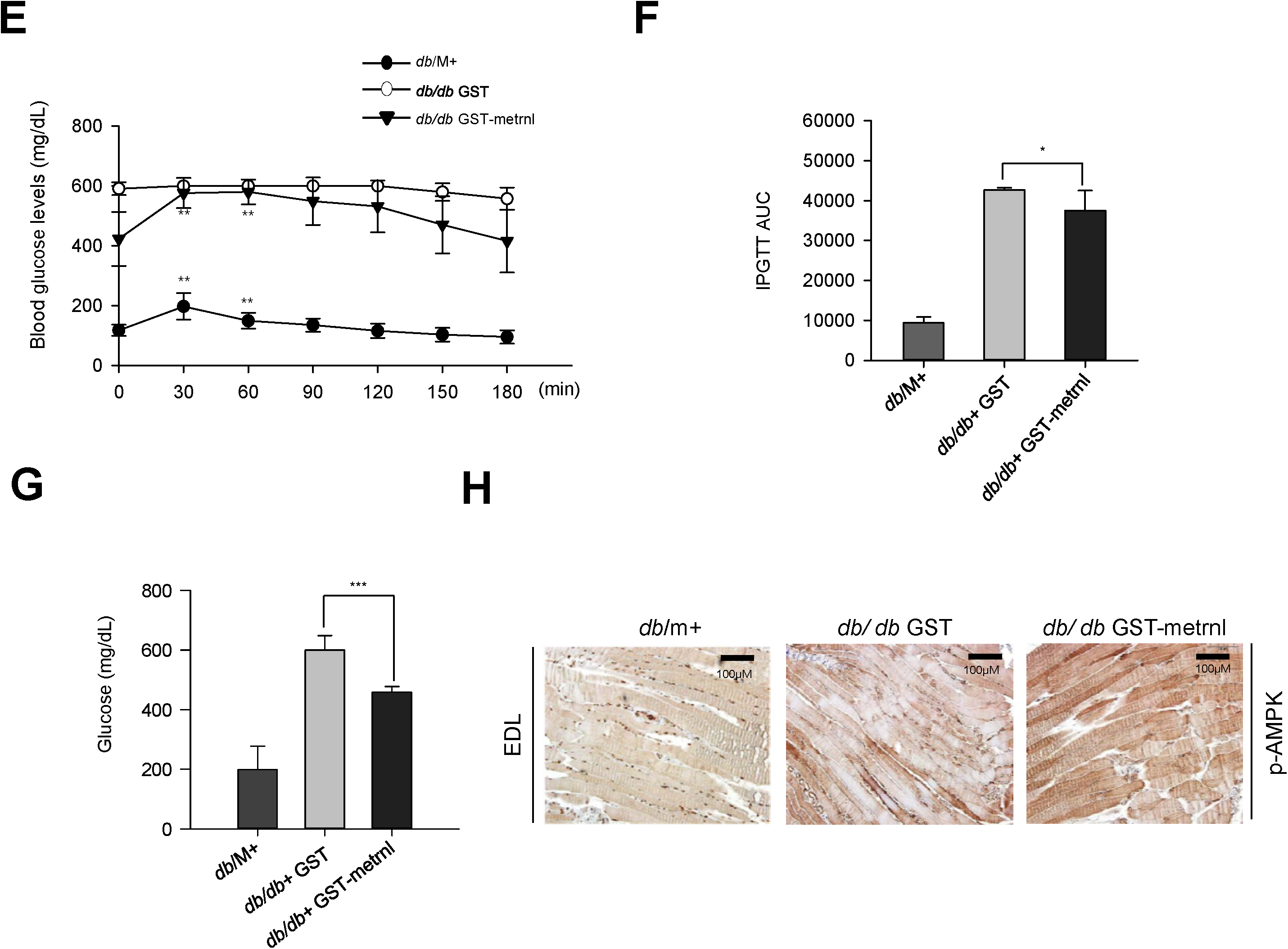
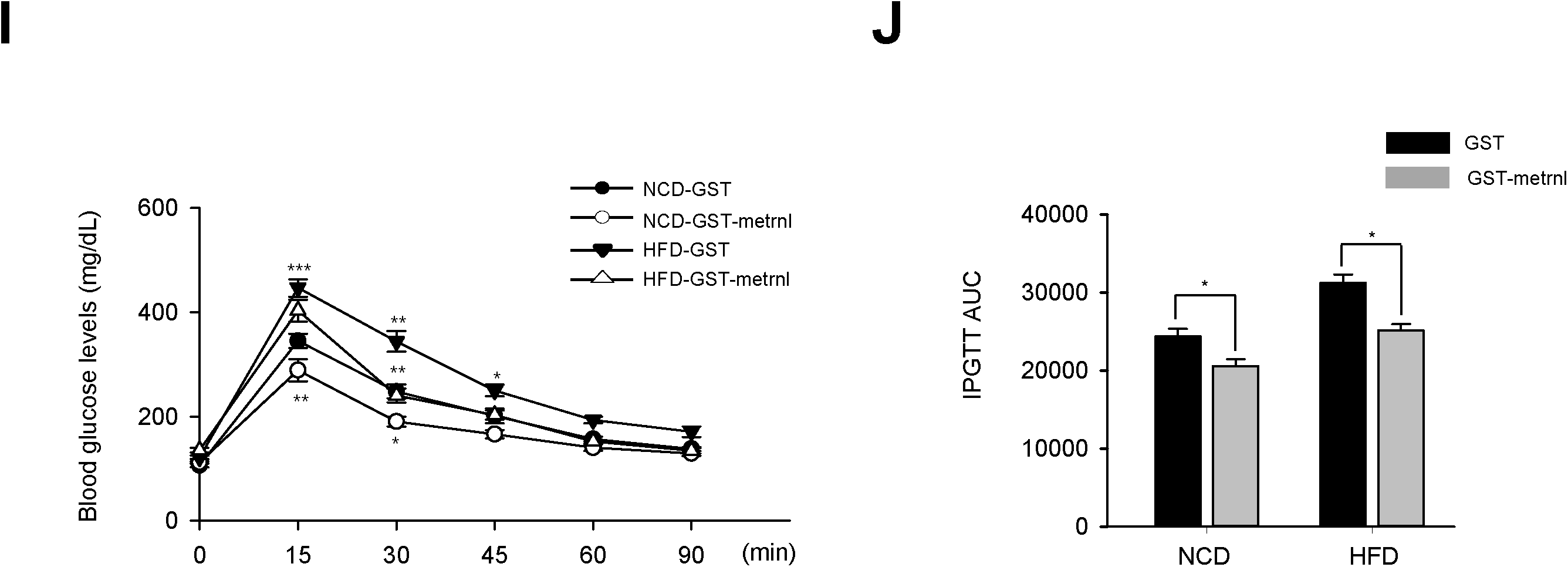
Metrnl improves glucose tolerance in diabetic (db/db) mice and mice with a normal chow diet (NCD) and high-fat diet (HFD). **a**. Recombinant metrnl-GST and GST protein were isolated using glutathione beads. The beads were washed three times with washing buffer, eluted, and analyzed by SDS-PAGE and subsequent Coomassie staining. **b**. C2C12 cells were treated with recombinant metrnl-GST. Cell lysates were analyzed with Western blotting using an antibody against phospho-AMPKα (Thr^172^), with AMPKα2 and β-actin as controls. **c**. Blood glucose concentrations and **d**. Area under the curve (AUC) results for a glucose tolerance test (GTT) in C57BL/6 mice. **e**. Blood glucose concentrations and **f**. Area under the curve (AUC) results for a glucose tolerance test (GTT) in db/M^+^, db/db + GST, and db/db + metrnl-GST mice. **g**. Blood glucose values for db/M^+^, db/db + GST, and db/db + metrnl-GST mice. **h**. Histochemistry of quadriceps muscles from db/M^+^, db/db + GST, and db/db + metrnl-GST mice. Each paraffin section was stained with anti-phospho-AMPKα2 antibodies. **i**. Blood glucose concentrations and **j**. Area under the curve (AUC) results for a glucose tolerance test (GTT) in mice fed a high-fat diet (HFD) and mice fed a normal chow diet (NCD), after injection with recombinant metrnl-GST or GST protein. Groups were compared using analysis of variance (ANOVA) with Duncan’s multiple range test. Results are displayed as the mean ± SD from three experiments. \**P* < 0.05 and \*\**P* < 0.01 vs. control, as indicated.

### AMPK β1β2 muscle-specific null mice showed no improvement in glucose tolerance after injection with metrnl

The AMP-activated protein kinase (AMPK) β1 and β2 subunits are required for assembling AMPK heterotrimers and are important for regulating enzyme activity. Mice lacking both the β1 and β2 isoforms in skeletal muscle (β1β2M-KO) had a drastically impaired capacity for treadmill running and contraction-stimulated glucose uptake^37^. To confirm the role of AMPK in observed effects on glucose metabolism, we administered recombinant metrnl into AMPK β1β2 muscle-specific null mice. Wild-type mice exhibited improved glucose tolerance upon metrnl intraperitoneal injection, which was not shown in AMPK β1β2 muscle-specific null mice (Fig. 9a and b). To further characterize the role of AMPK *in vivo*, we isolated EDL muscles from both WT mice and AMPK β1β2M-KO mice and measured glucose uptake ability. Recombinant metrnl increased glucose uptake in the EDL muscles of WT mice, but not in the EDL muscles of AMPK β1β2M-KO mice (Fig. 9c). These results clearly demonstrated that metrnl improves glucose tolerance via AMPK *in vivo*.

**Fig 9.**
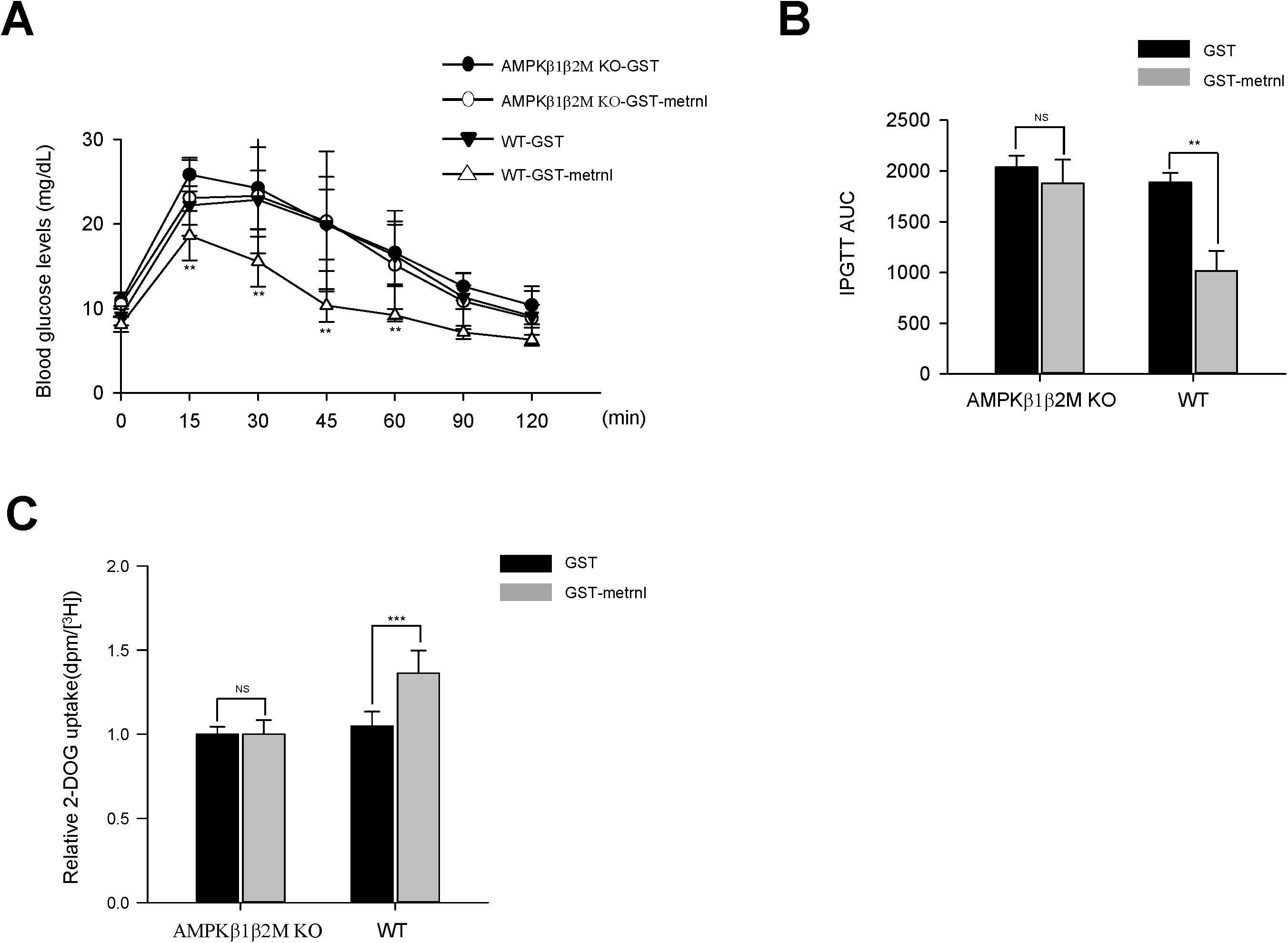
AMPK β1β2M-KO mice have reduced glucose tolerance and skeletal muscle glucose uptake after treatment with metrnl. **a**. Blood glucose concentrations and **b**. Area under the curve (AUC) results for a glucose tolerance test (GTT) in AMPK β1β2M-KO mice, after injection with recombinant metrnl-GST or GST protein. **c**. Extensor digitorum longus (EDL) tissues isolated from AMPK WT and AMPK β1β2M-KO mice were pre-incubated ± metrnl-GST - or GST alone (4µg/mL) and measured for 2-deoxyglucose (2DG) uptake. The data are presented as the relative 2-DG uptake (dpm/[H^3^] ± SD), based on 12-13 mice per group.

## Discussion

In this study, we found that metrnl stimulated glucose uptake and improved glucose tolerance via AMPK, both *in vitro* and *in vivo*. Circulating metrnl prompts energy expenditure and improves insulin sensitivity in mice^25^, but the underlying molecular mechanism is still not explored. Exercise improves glucose uptake through AMPK in an insulin-independent manner^35-40^. These studies point to a potential role for AMPK in exercise-mediated myokine function that warrants further investigation. The principal finding of the present study is that the CaMKK2-AMPK-HDAC5 axis plays an important role in metrnl-mediated glucose regulation in skeletal muscle.

AMPK is activated by two distinct mechanisms: an AMP-dependent pathway and a Ca^2+^-dependent pathway. LKB1 mediates AMPK activation via the AMP-dependent pathway, while CaMKK2 mediates activation via the Ca^2+^-dependent pathway^41^. Calcium is a molecular requirement for muscle contraction^42^. In a previous study, the metrnl gene was identified as a candidate for association with activator protein-1 (AP-1)^43^. Pax was also identified as a direct upstream transcription factor of metrnl^44^. There were few reports of the interaction between Pax and calcium, even though calcium is known as a critical factor for AP-1 activation^45^. In our study, metrnl increased intracellular calcium concentrations (Fig.3a). This implied that contraction may lead to metrnl gene transcription via the release of calcium from the lateral sacs of the sarcoplasmic reticulum. Histone deacetylases are enzymes that regulate gene expression by altering chromatin structure. Thus far, HDAC has been regarded as a regulator of skeletal muscle glucose metabolism. HDAC inhibition improves insulin sensitivity^46^ and diabetes mellitus^47^. In the present study, metrnl phosphorylated HDAC5, and phosphorylated HDAC5 became bound to 14-3-3 proteins. These complexes then translocated to the cytosol. Inhibition of AMPK blocked metrnl-mediated HDAC5 phosphorylation (Fig. 5c). These results imply that metrnl increases GLUT4 transcription via the phosphorylation of HDAC5 by AMPK. This is the first example in which a muscle-secreted molecule (metrnl) regulates GLUT4 through HDAC5. TBC1D1 and TBC1D4(AS160) are Rab-GTPase-activating proteins (Rab-GAPs)^48^. Phosphorylation of these protein stimulates GLUT4 mobilization to the plasma membrane^49^. TBC1D1 contains an AMPK-phosphorylation site (Ser237)^50-52^. This site apparently contributes to the unique regulatory aspects of TBC1D1. Furthermore, TBC1D4 phosphorylation at residue Thr642 is a key requirement for insulin-dependent glucose uptake by skeletal muscle^53^. In our study, metrnl increased the phosphorylation of TBC1D1 (Ser237) (Fig. 6a, b), but did not alter the phosphorylation of TBC1D4 (Thr642) (data not shown). Nevertheless, metrnl still promoted the translocation of GLUT4 to the cell membrane. These data demonstrated that metrnl stimulates GLUT4 translocation specifically via AMPK-TBC1D1, independent of insulin signaling.

In the present study, we found that metrnl improves glucose tolerance in both obese and type 2 diabetic mouse models, but without dramatic weight loss. In contrast, Rao et al. reported that circulating metrnl does lead to weigh loss^25^. In our case, the recombinant GST-metrnl protein was intraperitoneally injected 3 times per week for 15 weeks at the beginning of high-fat diet feeding to investigate the role of metrnl in the initiation of obesity. It slightly increased the body weight in both normal chow diet and high fat diet groups (data not shown), but improved glucose tolerance in the all groups. On the other hands, mice fed a high-fat diet 60% (HFD) for 20 weeks were injected daily with saline or metrnl-Fc protein for 7 days by Rao et al. Additional studies are needed to clarify the role of metrnl on the body weight. To further confirm the role of AMPKα in mediating anti-diabetes effects via metrnl, we examined the effect of metrnl in AMPK β1β2 muscle-specific null mice. Metrnl did not improve glucose uptake or tolerance in AMPK β1β2 muscle-specific null mice compared with wild-type mice (Fig. 9a and c). These results demonstrated that metrnl improves glucose tolerance *in vivo* through the AMPK signaling pathway.

Very recently, the association of circulating levels of meteorin with cardiometabolic risk factors and type 2 diabetes (T2DM) was inconsistently reported. Chung et al. Reported that the levels of metrnl were higher in T2DM and circulating levels of metrnl were negatively associated with metabolic syndrome components such as blood pressure, total-and LDL cholesterol and fasting plasma glucose. In that study, they speculated that the observed high level of metrnl in diabetes might be a defensive response to counteract metabolic stress in dealing with hyperglycemia^54^. This is in line with our preliminary results that serum metrnl level was elevated in T2DM in a small subset of free-living elderly population (n=60, data not shown). On the other hand, another study^55^ reported that serum metrnl levels were significantly lower in patients who had been newly diagnosed T2DM, raising the question as to the duration of diabetes may influence the blood metrnl status and its metabolic function. In the future, further studies will be necessary to explain the confounding relationship between metrnl levels and type 2 diabetes.

In conclusion, we have demonstrated for the first time that exercise-induced metrnl has an anti-diabetes effect via the AMPKα signaling pathway. Moreover, these results provide evidence of the beneficial effects of exercise-mediated adipomyokines on glucose homeostasis. Further studies could offer more insight into the possible utility of metrnl for T2DM prevention and treatment.

## Methods

### Reagents

Antibodies against phospho-AMPKα(Thr^172^), AMPKα, AMPKα2, phospho-ACC (Ser^79^), ACC, and p-TBC1D1 (Ser^237^) were purchased from Merck Millipore (Darmstadt, Germany). Antibodies against GLUT4 and 14-3-3 were purchased from Abcam (Boston, MA, USA). Antibodies against p38 MAPK was purchased from Cell Signaling Technology (Danvers, MA, USA). The antibody against β-actin was purchased from Sigma-Aldrich (St. Louis, MO, USA). Antibodies against phospho-p38 MAPK, α-tubulin, GLUT4, and lamin B were purchased from Santa Cruz Biotechnology (Santa Cruz, CA, USA). Horseradish peroxidase (HRP)-conjugated goat anti-rabbit IgG and goat anti-mouse secondary antibodies were purchased from Enzo Life Sciences (Farmingdale, NY, USA). Metrnl was obtained from Cusabio (Wuhan, Hubei, China), and 1,2-bis (o-aminophenoxy) ethane-N,N,N’,N’-tetraacetic acid (BAPTA)-AM was purchased from Abcam. Compound C and STO-609 were obtained from Calbiochem (San Diego, CA, USA). Protein A-agarose beads were obtained from GE Healthcare (Piscataway, NJ, USA). The fluorescent Ca^2+^ indicator Fluo-3 AM and Hoechst 33342 were obtained from Invitrogen (Leiden, Netherlands).

### Cell culture

Mouse myoblast C2C12 cells and Rat myoblasts L6 (ATCC; Rockville, MD, USA) were cultured in Dulbecco’s modified Eagle medium (DMEM; Invitrogen) supplemented with 10% fetal bovine serum (FBS) and 1% antibiotics at 37°C in a 5% CO_2_ atmosphere.

### Ca2+ measurement

Cells were treated with 5 μM Fluo-3 AM in regular culture medium at 37°C for 30 minutes. Cells were then washed and incubated for 15 min in regular medium (without Fluo-3 AM) to complete the de-esterification process. The cells were treated with metrnl, and the culture plates were placed on a temperature-controlled confocal microscope (Zeiss LSM 700 Meta; Zeiss, Oberkochen, Germany) at 200X magnification. The excitation and emission wavelengths for signal detection were 488 nm and 515 nm, respectively.

### Immunoblot analyses

Following the various experimental manipulations, the culture medium was removed, and the cells were washed twice with ice-cold phosphate-buffered saline (PBS). They were then lysed with 70 μL of lysis buffer [50 mM Tri-HCl (pH 7.4), 1% Triton X-100, 0.25% sodium deoxycholate, 150 mM EDTA, 1 mM sodium orthovanadate (Na_3_VO_4_), 1 mM NaF, and 1 mM phenylmethylsulfonyl fluoride (PMSF)]. Samples were sonicated, centrifuged for 20 minutes, and heated at 95°C for 5 minutes. Proteins were resolved on 10% SDS-PAGE gels and transferred to nitrocellulose membranes. Membranes were blocked in Tris-buffered saline with 0.1% Tween 20 (TBS-T buffer) and 5% dry milk (w/v) for 1 h, then washed 3 times in TBS-T. Membranes were incubated overnight at 4°C with primary antibodies and probed with HRP-conjugated secondary antibodies for 1 h. The blots were visualized via chemiluminescence using the ECL detection system (Amersham International PLC, Buckinghamshire, UK).

### Silencing AMPKα and TBC1D1

Transient transfections were performed with Lipofectamine 2000 (Invitrogen, Carlsbad, CA, USA) according to the manufacturer’s protocol. Briefly, 5 μL of siRNA targeted to mouse TBC1D1 (L-040360-00, On-TARGET plus SMART pool oligonucleotide, NM_001289514; Dharmacon, Lafayette, CO, USA) contained the following sequences: CGACGAAUACCUUCGCCAAA, CUGCACAAGCUCUGCGAAA, GUGAGGAGAGGCGUUCAA, and CAAGAAACGAGCAGCGAGA and mouse AMPKα2 (L-NM_100623,, On-TARGET plus SMART pool oligonucleotide, NM_023991; Dharmacon, Lafayette, CO, USA) contained the following sequences: UAUCUCAACCGUUCUAUUG, GUGAAUGCAUACCAUCUUC, GACUGGCAAUUACGUGAAA, andGAAGAUUCGCAGUUUAGAU. Similarly, 5 μL of siRNA targeted to mouse non-targeting pool (L-001810-10, On-TARGET plus SMART pool oligonucleotide; Dharmacon) contained the following sequences: UGGUUUACAUGUCGACUAA, UGGUUUACAUGUUGUGUGA, UGGUUUACAUGUUUUCUGAGUGA, andUGGUUUACAUGUUUUCCUA. For each experiment, Lipofectamine 2000 (5 μL) was diluted in 95 μL of reduced-serum medium (Opti-MEM; Invitrogen), then mixed with siRNA. The mixtures were incubated for 15 min before being added dropwise to culture wells containing 800 μL of Opti-MEM, to achieve a final siRNA concentration of 50 nM.

### Reverse transcription-polymerase chain reaction (RT-PCR) and real-time PCR

RT-PCR was performed at 55°C for 20 minutes using a Thermoscript II one-step RT-PCR Kit (Invitrogen). cDNA amplification was carried out using a Gene Amp System 9700 thermocycler (Applied Biosystems, Warrington, UK). The reverse transcriptase was heat-inactivated in the first step of the PCR reaction (95°C for 10 minutes). The following primers were used for amplification: metrnl, 5′-AAGCCTTTCAGGGACTCCTC-3′ (sense) and 5′-CCCTGGTCGTACTCCACACT-3′ (antisense); β-actin, 5′-ATTTGGTCGTATTGGGCGCCTGGTCACC-3′ (sense) and 5′-GAAGATGGTGATGGGATTTC-3′ (antisense); GLUT4 for real time PCR, 5′-AGCTGGTGTGGTCAATACGG-3′ (sense) and 5′-AACAGATGGAGTGTCCGTCG-3′ (antisense); GLUT4 for ChIP, 5′-CTTCGACCTTTCAGGGGGAC-3′ (sense) and 5′-GAACAAAAGGCTCTTCCCGC-3′ (antisense). Amplification steps were as follows: 32 cycles at 95°C for 15 seconds, 58°C (β-actin) or 55°C (GLUT4 and metrnl) for 30 seconds, and 72°C for 30 seconds, followed by 10 minutes at 72°C. A 10 µL sample of each reaction was analyzed by agarose gel electrophoresis. In case of real-time PCR, the relative amount of target genes was performed by measuring the threshold cycle (CT) values of target genes and β-actin. The relative amount of target genes was normalized by the amount of β-actin used as the internal control in the same sample and described as the ratio of each target gene/β-actin.

### Immunodetection of Myc-GLUT4

Cell surface expression of Myc-GLUT4 was quantified using an antibody-coupled colorimetric absorbance assay, as described previously^18^. Following stimulation, differentiated L6 myotubes stably expressing Myc-GLUT4 were incubated with a polyclonal anti-Myc antibody (1:1000) for 60 minutes and incubated with an HRP-conjugated goat anti-rabbit IgG (1:1000) for 1 h. Cells were washed six times with PBS and incubated in 1 mL OPD reagent (0.4 mg/mL) for 30 minutes. Absorbance of the supernatant was measured at 492 nm.

### Glucose uptake

Two days after cells reached confluence, L6 myotube differentiation was induced by incubation in DMEM supplemented with 2% FBS for 6–7 days; the growth medium was changed every 2 days. Cells were washed twice with PBS, then starved in serum-free low-glucose DMEM for 3 hours. Cells were incubated with KRB (20 mM HEPES [pH 7.4], 130 mM NaCl, 1.4 mM KCl, 1 mM CaCl_2_, 1.2 mM MgSO_4_, and 1.2 mM KH_2_PO_4_), then incubated with test compounds in the same buffer at 37°C. The uptake assay was initiated by adding 2-deoxy-D-(H^3^)-glucose (2-DG) to each well and incubating at 37°C for 15 minutes. The reaction was terminated by washing with ice-cold PBS. Cells were lysed in 10% SDS. An aliquot of the cell lysate was removed for protein quantitation by the method of Bradford assay. The amount of [H^3^]-2-deoxyglucose uptake was determined (in triplicate) by scintillation counting.

### Immunoprecipitation

Cellular proteins (1 mg) were mixed with 1 μg of anti-14-3-3 (rabbit monoclonal antibody) or anti-IgG (normal rabbit antibody), and incubated at 4°C for 24 h. Immune complexes were captured with protein A-Sepharose (Amersham, Uppsala, Sweden) for an additional 3 h. Precipitated immune complexes were washed 3 times with wash buffer [25 mM HEPES, 5 mM EDTA, 1% Triton X-100, 50 mM NaF, 150 mM NaCl, 10 mM PMSF, 1 μM leupeptin, 1 μM pepstatin, and 1 μM aprotinin (pH 7.2)]. The washed samples were resuspended in SDS sample buffer [125 mM Tris-HCl (pH 6.8), 20% (v/v) glycerol, 4% (w/v) SDS, 100 mM dithiothreitol, and 0.1% (w/v) bromophenol blue], and heated at 100°C for 5 min.

### Isolation of the plasma membrane fraction

C2C12 mouse myoblast cells (2×10^7^) were plated in 10 cm treated cell culture dishes, and the growth medium was changed to Opti-MEM for 6 h. After that, the cells were treated with metrnl (100ng/mL) for 3 h, or with 100 nM insulin for 30 min. The supernatants were removed, and cells were washed three times with ice-cold PBS. The plasma membrane was extracted and purified by a plasma membrane protein extraction kit (ab65400; Abcam, Boston, MA, USA), according to the manufacturer’s instructions.

### Chromatin immunoprecipitation assay

A ChIP assay was performed using a kit (Cell Signaling Technology, MA, USA), according to the manufacturer’s instructions. C2C12 myoblasts were treated with metrnl, and DNA-protein complexes were cross-linked using 1% formaldehyde for 15 min, followed by quenching with 125 mmol/L glycine. Cross-linked chromatin samples were isolated from the cell lysate using nuclease digestion. HDAC5 was immunoprecipitated using an HDAC5 antibody (Novus Biologicals; Littleton, CO, USA), or an unrelated antibody (normal rabbit IgG) for a control. DNA was extracted. For quantitative PCR assays, ChIP DNA was amplified using primers designed to amplify the GLUT4 promoter containing the MEF2 binding site, as follows: 5′-CTT CGA CCT TTC AGG GGG AC-3′ (forward) and 5′-GAA CAA AAG GCT CTT CCC GC-3′ (reverse). Each reaction utilized Power SYBR green PCR Master Mix from Applied Biosystems (Foster City, CA, USA). Values represent enrichment over the IgG negative control, using the threshold cycle (2^-ΔΔCT^) method.

### Fractionation of nuclear and cytosolic proteins

A plasma membrane protein extraction kit (Thermo Scientific 78833; Rockford, IL, USA) was used to obtain plasma membrane fractions from cells, according to the manufacturer’s protocol. Briefly, exponentially growing cultures of C2C12 cells were collected using a cell scraper and washed at least twice with cold PBS (pH 7.4) by resuspension and centrifugation. The cells were then mixed with cytoplasmic extraction reagent (CER) I. The cell pellet was gently resuspended, incubated for 10 min on ice, and then ice-cold CER II was added. The tube was vortexed for 5 s on the highest setting, incubated on ice for 1 min, and centrifuged at 13,000 rpm for 15 min. The cell supernatant contained cytoplasmic proteins. To extract nuclear proteins, the pellet was suspended in ice-cold NER, vortexed, placed on ice for 40 minutes, and centrifuged at 13,000 rpm for 10 min. The supernatant (nuclear extract) fraction was immediately transferred to a tube.

### Immunocytochemistry

Cells were fixed with 4% paraformaldehyde (PFA)/PBS at room temperature for 15?minutes. After blocking with 3% bovine serum albumin (BSA) at room temperature for 30?minutes, fixed cells were incubated with primary anti-p-HDAC5 (1:500, gtx50238; GeneTex), primary anti-p-TBC1D1 (1:500, 07-2268; Merck Millipore), or primary anti-14-3-3 (1:500, ab6081; Abcam) antibodies in primary antibody diluent (PBS, 3% BSA, and 0.1% Triton-X100). This was followed by incubation at 4°C overnight. Cells were then washed with PBS, probed with a goat anti-rabbit Cy3 (red) or goat anti-mouse 488 (green) secondary antibody (Molecular Probes, Eugene, OR, USA), and washed 3 times with PBS at room temperature for 10?minutes. Images were obtained with a Zeiss confocal microscope.

### Plasmid construction of GLUT4-GFP and metrnl-GST

A mouse myc-DDK-tagged GLUT4 ORF clone (MR_208202) and a mouse metrnl ORT clone (MR_14497) were purchased from Origene Technologies, Inc. (Rockville, MD, USA). Plasmid DNA from these clones was amplified by PCR, using the following primers: GLUT4 forward primer, 5’-CGCGGGCCCGGGATCC ATG CCT TCG GGT TTC CAG CAG-3’; GLUT4 reverse primer 5’-G AGC TCG CAA ACA GAG CTG AAC TAG-3; metrnl forward primer, 5’-GGT TCC GCG TGG ATC CCA GTA CTC CAG CGA CCT G-3’; and metrnl reverse primer 5’-GAT GCG GCC GCT CGA GCT CCA TAT TGA TTT CAC A-3’. BamH1-and Sac1-digested products of GLUT4, and BamH1-and Xho1-digested products of metrnl were respectively ligated into a linearized pEGFP-C1 vector (Clontech; Palo Alto, CA, USA) or a pGEX4-1 vector (GE Healthcare; Boston, MA, USA). All constructs were verified by direct sequencing. Nuclei were stained with Hoechst 33342 dye for 30 min at 25°C. Confocal images were obtained using a Zeiss confocal microscope (LSM700) and analyzed with the Zeiss LSM image browser software (Carl Zeiss, Oberkochen, Germany).

### Confocal microscopy

C2C12 cells expressing GLUT4-GFP were fixed with 4% PFA/PBS at room temperature for 15?minutes. After blocking with 3% BSA at room temperature for 30?min, nuclei were stained with Hoechst 33342 dye for 30 min at 25°C. Confocal images were obtained using a Zeiss confocal microscope (LSM700) and analyzed with the Zeiss LSM image browser software (Carl Zeiss, Oberkochen, Germany).

### Immunohistochemistry

Representative blocks of paraffin-embedded tissues were cut at 4 μm thickness, dewaxed, and rehydrated. Briefly, sections were deparaffinized, rehydrated, and washed in PBS. To block nonspecific binding, sections were incubated in 4% BSA+ dextran for 1 h at 4°C. Sections were incubated with anti-AMPKα2 and anti-GLUT4 antibodies at a dilution of 1:200 (in 1% BSA and 0.1% Nonidet P-40 in PBS) overnight at 4°C. The Vectastain ABC kit (Vector Labs Burlingame, CA, USA) was used for the avidin-biotin complex (ABC) method, according to the manufacturer’s instructions. Peroxidase activity was visualized with 3,3′-diaminobenzidine (Darko, Carpinteria, CA, USA). The sections were lightly counterstained with hematoxylin, dehydrated through an ethanol series to xylene, and mounted.

### Protein purification of metrnl-GST and GST, and administration in mice

GST fusion proteins, including GST only and GST-metrnl, were expressed in *E. coli* and purified using glutathione agarose beads, according to manufacturer’s instructions (GE Healthcare). The purity and integrity of the fusion proteins were analyzed by SDS-PAGE and Coomassie Blue staining. Diabetic (db/db) and HFD mice were administered recombinant metrnl-GST or GST only at 3 mg/kg (through intraperitoneal injection) for 4 weeks.

### Mouse and human metrnl sandwich ELISA assay

Immediately after exercise, blood was collected from control and exercised mice. Blood samples were centrifuged at 3,000 rpm for 10 min at 4°C. The serum metrnl concentration was measured using a mouse metrnl ELISA Kit (Cusabio), according to the manufacturer’s instructions.

### *Ex vivo* glucose uptake

Primary myoblasts were obtained from the forelimbs and hindlimbs of 5-day-old littermate pups (n = 3-4). Dissected and minced muscle was enzymatically disaggregated at 37°C in 4 mL PBS containing 1.5 U/mL dispase II and 1.4 U/mL collagenase D (Roche, Penzberg, Germany). Samples were sheared by mixing with a 10 mL pipette every 5 minutes for 20 minutes. Cells were filtered through 70 µm mesh (BD, Seoul, Korea) and collected by pelleting at 1,000 rpm for 5 minutes. The cell pellet was dissociated in 10 mL F10 medium (Invitrogen) supplemented with 10 ng/mL basic fibroblast growth factor (PeproTech, NJ, USA) and 10% Cosmic Calf Serum (Hyclone, Logan, UT, USA). Cells were pre-plated twice onto non-collagen-coated plates for 1 h each, to deplete the fibroblasts. After cells reached confluence, differentiation was induced by incubation in DMEM supplemented with 2% FBS for 2 days. Cells were washed twice with PBS, then starved in serum-free, low-glucose DMEM for 3 h. Cells were incubated with KRB, then incubated with the indicated compounds at 37°C. The uptake assay was initiated by adding 2-deoxy-D(H3)-glucose (2-DG) to each well, and incubating at 37°C for 15 min. The reaction was terminated by washing with ice-cold PBS. The cells were lysed in 10% SDS and mixed with a scintillation cocktail to measure radioactivity. The experiment was approved by the Korea University Institutional Animal Care and Use Committee (IACUC, KOREA-2016-0252). It was performed in accordance with the guidelines and regulations of the IACUC. The glucose uptake experiment in Figure 9C was conducted as previously described^56^ in the presence of GST-metrnl or GST alone (4µ g/mL).

### Electric pulse stimulation

Electric pulse stimulation (EPS) was applied to cells using a C-Pace EP culture pacer (IonOptix, MA, USA), which is a multi-channel stimulator designed for chronic stimulation of bulk quantities of cells in culture. This instrument emits bipolar pulses to culture media via immersed carbon electrodes in a C-dish. EPS was applied to differentiated L6 cells cultured under conditions of high-density micro-mass (2 × 10^5^ cells/mL) under electrical fields of 25 V/cm, with a duration of 5 m/s and a frequency of 1 Hz. At appropriate time points, cells were harvested in Trizol (Invitrogen) for PCR analysis, or in lysis buffer for Western blotting.

### Chronic forced treadmill running procedure

Eight-week-old, specific-pathogen-free, male BALB/C mice (Dae-Han Biolink Company, Chungcheongbuk-do, Korea) were maintained according to Korea University College of Medicine research requirements. All procedures were approved by the Committee on Animal Research at the Korea University College of Medicine (KOREA-2016-0252). The animals were fed chow and water ad libitum and were acclimatized to a 12 h light cycle (lights on between 0600 and 1800 h) for a period of 1 week before experimental manipulation. Mice were divided into two groups of which one group received treadmill training (20cm diameter and 5 cm width; Shandong Yiyuan Technology Development Co., Shandong, Binzhou, China) for three weeks. The forced exercise was performed at a velocity of 10 m/min for 60 min. This forced exercise was administered 5 days/week. Mice that failed to run were given a 2.0 mA electric shock, and the running equipment was stopped for 30 s when mice reached exhaustion. The number of electric shocks and running times during exercise were automatically recorded by the running wheel equipment. After the final exercise session on day 21, animals were anesthetized with Zoletil (Virvac Laboratories, Carros, France) by intraperitoneal injection. Blood samples were harvested by cardiac puncture into tubes containing heparin solution. Samples were centrifuged at 2000 × *g* for 10 min to obtain plasma. The plasma was used for ELISA testing (metrnl ELISA kit; Cusabio, Wuhan, Hubei, China).

### Animals and experimental design

For the diet-induced obesity experiments, 50 specific-pathogen-free C57BL/6N male mice (7 weeks old, 22–24g) were obtained from Koatech (Gyeonggi-do, Korea). Mice were then fed a high-fat diet (HFD) or a normal chow diet (NCD). Seven-week-old male db/m and db/db mice (C57BL KSJ M^+^/lepR^-/-^) were supplied by Central Lab Animal, Inc. (Seoul, Korea) for the diabetic mouse experiments. All experimental animals were maintained according to Korea University College of Medicine research requirements, and all procedures were approved by the Committee on Animal Research at the Korea University College of Medicine (KOREA-2016-0252). For the obese mice, eight-week-old C57BL/6N males were divided randomly into four groups of ten animals per group. The first and second groups were then fed HFD and injected with recombinant GST or recombinant metrnl-GST for 8 weeks. The third and fourth groups were fed NCD and injected with GST or recombinant metrnl-GST for 8 weeks. For the diabetic mice, eight-week-old db/db males were divided randomly into two groups with ten animals per group. The db/m group was the positive control. The first group of db/db mice was injected with recombinant GST, and the second group was injected with recombinant metrnl-GST. These injections were given 3 times per week for 8 weeks. After the final treatment with recombinant protein, glucose tolerance tests (GTTs) were performed. All experimental animals fasted for ∼16 h before 20% glucose (2 g/kg) was injected intraperitoneally. Blood glucose levels were measured before injection and 15, 30, 60, and 120 min after injection. Blood glucose concentrations were measured using an Accu-Check glucometer (Roche).

### Data analysis

All data are presented as means±standard error of the mean. Differences in treatment groups were tested using one-way ANOVA with Tukey’s post-tests or Student’s t-test. Differences with p < 0.05 were considered statistically significant. Statistical analysis was performed Sigma Plot 12.0. for Windows (Systat Software Inc., London, UK).

## Acknowledgements

This study was supported by the National Research Foundation of Korea, which is funded by the Korean government (NRF-2016R1610722).

## Author Contributions

The experiments were conceived and designed by Kim Hyeon Soo and Lee Jeong Ok. The experiments were performed by Lee Jeong Ok, Jeong Ah Han, Min Ju Kang, Ilhyeok Seo, Shin Ae Kim, Yong Woo Lee, Lisa Murray-Segal, Kevin R.W. Ngoei, and Naomi X.Y. Ling. Reagents and/or analysis tools were contributed by Min-Jeong Shin, Hye Jeong Lee, Eun-Soo Lee, Hong min Kim, Choon Hee Chung, Kevin R.W. Ngoei, Naomi X.Y. Ling, Jonathan S. Oakhill, Sandra Galic, Jiyoung Moon, Kyoung Min Kim, and Soo Lim. The following authors reviewed and contributed to revisions of the manuscript: Kevin R.W. Ngoei, Naomi X.Y. Ling, Jonathan S. Oakhill, Sandra Galic, Lisa Murray-Segal, and Bruce E. Kemp.

## Competing Interests

The authors declare no competing interests.

## References

1. Booth, F. W., Roberts, C. K. & Laye, M. J. Lack of exercise is a major cause of chronic diseases. Compr. Physiol. 2, 1143–1211 (2012).

2. Hoffmann, C. & Weigert, C. Skeletal Muscle as an Endocrine Organ: The Role of Myokines in Exercise Adaptations. Cold Spring Harb. Perspect. Med. 7(2017).

3. Pedersen, B. K., Akerstrom, T. C., Nielsen, A. R. & Fischer, C. P. Role of myokines in exercise and metabolism. J. Appl. Physiol. (1985) 103, 1093–1098 (2007).

4. Pedersen, B. K. The diseasome of physical inactivity--and the role of myokines in muscle--fat cross talk. J. Physiol. (Lond.) 587, 5559–5568 (2009).

5. Pedersen, B. K. Muscles and their myokines. J. Exp. Biol. 214, 337–346 (2011).

6. Busquets, S., Figueras, M., Almendro, V., Lopez-Soriano, F. J. & Argiles, J. M. Interleukin-15 increases glucose uptake in skeletal muscle. An antidiabetogenic effect of the cytokine. Biochim. Biophys. Acta 1760, 1613–1617 (2006).

7. Barra, N. G., Chew, M. V., Holloway, A. C. & Ashkar, A. A. Interleukin-15 treatment improves glucose homeostasis and insulin sensitivity in obese mice. Diabetes Obes. Metab. 14, 190–193 (2012).

8. Alvarez, B. et al. Effects of interleukin-15 (IL-15) on adipose tissue mass in rodent obesity models: evidence for direct IL-15 action on adipose tissue. Biochim. Biophys. Acta 1570, 33–37 (2002).

9. Pedersen, B. K. & Febbraio, M. A. Muscle as an endocrine organ: focus on muscle-derived interleukin-6. Physiol. Rev. 88, 1379–1406 (2008).

10. Serrano, A. L., Baeza-Raja, B., Perdiguero, E., Jardi, M. & Munoz-Canoves, P. Interleukin-6 is an essential regulator of satellite cell-mediated skeletal muscle hypertrophy. Cell Metab. 7, 33–44 (2008).

11. Broholm, C. & Pedersen, B. K. Leukaemia inhibitory factor--an exercise-induced myokine. Exerc. Immunol. Rev. 16, 77–85 (2010).

12. Ouchi, N., Parker, J. L., Lugus, J. J. & Walsh, K. Adipokines in inflammation and metabolic disease. Nat. Rev. Immunol. 11, 85–97 (2011).

13. Bluher, M. Clinical relevance of adipokines. Diabetes Metab. J. 36, 317–327 (2012).

14. Wang, P. et al. Involvement of leptin receptor long isoform (LepRb)-STAT3 signaling pathway in brain fat mass-and obesity-associated (FTO) downregulation during energy restriction. Mol. Med. 17, 523–532 (2011).

15. Miao, C. Y. Introduction: Adipokines and cardiovascular disease. Clin. Exp. Pharmacol. Physiol. 38, 860–863 (2011).

16. Li, Z. Y., Wang, P. & Miao, C. Y. Adipokines in inflammation, insulin resistance and cardiovascular disease. Clin. Exp. Pharmacol. Physiol. 38, 888–896 (2011).

17. Lee, H. J. et al. Irisin, a Novel Myokine, Regulates Glucose Uptake in Skeletal Muscle Cells via AMPK. Mol. Endocrinol. 29, 873–881 (2015).

18. Lee, H. J. et al. Kalirin, a GEF for Rac1, plays an important role in FSTL-1-mediated glucose uptake in skeletal muscle cells. Cell Signal. 29, 150–157 (2017).

19. Lee, J. O. et al. Resistin, a fat-derived secretory factor, promotes metastasis of MDA-MB-231 human breast cancer cells through ERM activation. Sci. Rep. 6, 18923 (2016).

20. Lee, J. O. et al. Visfatin, a novel adipokine, stimulates glucose uptake through the Ca2+-dependent AMPK-p38 MAPK pathway in C2C12 skeletal muscle cells. J. Mol. Endocrinol. 54, 251–262 (2015).

21. Görgens, S. W., Eckardt, K., Jensen, J., Drevon, C. A. & Eckel, J. Exercise and Regulation of Adipokine and Myokine Production. Prog. Mol. Biol. Transl. Sci. 135, 313–336 (2015).

22. Li, Z. Y. et al. Subfatin is a novel adipokine and unlike Meteorin in adipose and brain expression. CNS Neurosci. Ther. 20, 344–354 (2014).

23. Zheng, S. L., Li, Z. Y., Song, J., Liu, J. M. & Miao, C. Y. Metrnl: a secreted protein with new emerging functions. Acta Pharmacol. Sin. 37, 571–579 (2016).

24. Ushach, I. et al. METEORIN-LIKE is a cytokine associated with barrier tissues and alternatively activated macrophages. Clin. Immunol. 156, 119–127 (2015).

25. Rao, R. R. et al. Meteorin-like is a hormone that regulates immune-adipose interactions to increase beige fat thermogenesis. Cell 157, 1279–1291 (2014).

26. Zhang, B. B., Zhou, G. & Li, C. AMPK: an emerging drug target for diabetes and the metabolic syndrome. Cell Metab. 9, 407–416 (2009).

27. Long, Y. C. & Zierath, J. R. AMP-activated protein kinase signaling in metabolic regulation. J. Clin. Invest. 116, 1776–1783 (2006).

28. Crane, J. D. et al. Exercise-stimulated interleukin-15 is controlled by AMPK and regulates skin metabolism and aging. Aging Cell 14, 625–634 (2015).

29. Sarabia V, Ramlal T, Klip A. Glucose uptake in human and animal muscle cells in culture. Biochemistry and Cell Biology 68, 536–542 (1990).

30. Thomas E. Jensen, Yeliz Angin, Lykke Sylow & Erik A. Richter. Is contraction-stimulated glucose transport feedforward regulated by Ca2+?. Experimental Physiology 99, 1562–1568 (2014).

31. McFalls, E. O. et al. Activation of p38 MAPK and increased glucose transport in chronic hibernating swine myocardium. Am. J. Physiol. Heart Circ. Physiol. 287, H1328–1334 (2004). J. Biol. Chem 19, 13824–13829 (1993).

32. Watson, R. T. & Pessin, J. E. Bridging the GAP between insulin signaling and GLUT4 translocation. Trends Biochem. Sci. 31, 215–222 (2006).

33. P D Neufer, J O Carey & G L Dohm. Transcriptional regulation of the gene for glucose transporter GLUT4 in skeletal muscle. Effects of diabetes and fasting.

34. McKinsey, T. A., Zhang, C. L., Lu, J. & Olson, E. N. Signal-dependent nuclear export of a histone deacetylase regulates muscle differentiation. Nature 408, 106–111 (2000).

35. Grozinger, C. M. & Schreiber, S. L. Regulation of histone deacetylase 4 and 5 and transcriptional activity by 14-3-3-dependent cellular localization. Proc. Natl. Acad. Sci. U. S. A. 97, 7835–7840 (2000).

36. Zisman, A. et al. Targeted disruption of the glucose transporter 4 selectively in muscle causes insulin resistance and glucose intolerance. Nat. Med. 6, 924–928 (2000).

37. Chavez, J. A., Roach, W. G., Keller, S. R., Lane, W. S. & Lienhard, G. E. Inhibition of GLUT4 translocation by Tbc1d1, a Rab GTPase-activating protein abundant in skeletal muscle, is partially relieved by AMP-activated protein kinase activation. J. Biol. Chem. 283, 9187–9195 (2008).

38. O’Neill, H. M. et al. AMP-activated protein kinase (AMPK) beta1beta2 muscle null mice reveal an essential role for AMPK in maintaining mitochondrial content and glucose uptake during exercise. Proc. Natl. Acad. Sci. U. S. A. 108, 16092–16097 (2011).

39. Schrauwen, P. & van Marken Lichtenbelt, W. D. Combatting type 2 diabetes by turning up the heat. Diabetologia 59, 2269–2279 (2016).

40. Goodyear, L. J. AMP-activated protein kinase: a critical signaling intermediary for exercise-stimulated glucose transport? Exerc. Sport Sci. Rev. 28, 113–116 (2000).

41. Ost, M., Coleman, V., Kasch, J. & Klaus, S. Regulation of myokine expression: Role of exercise and cellular stress. Free Radic. Biol. Med. 98, 78–89 (2016).

42. Huang, S. & Czech, M. P. The GLUT4 glucose transporter. Cell Metab. 5, 237–252 (2007).

43. Gong, W. et al. Meteorin-Like Shows Unique Expression Pattern in Bone and Its Overexpression Inhibits Osteoblast Differentiation. PLoS One 11, e0164446 (2016).

44. Ramialison, M. et al. Rapid identification of PAX2/5/8 direct downstream targets in the otic vesicle by combinatorial use of bioinformatics tools. Genome Biol. 9, R145 (2008).

45. Monaco, S., Jahraus, B., Samstag, Y. & Bading, H. Nuclear calcium is required for human T cell activation. J. Cell Biol. 215, 231–243 (2016).

46. Ye, J. Improving insulin sensitivity with HDAC inhibitor. Diabetes 62, 685–687 (2013).

47. Christensen, D. P. et al. Histone deacetylase (HDAC) inhibition as a novel treatment for diabetes mellitus. Mol. Med. 17, 378–390 (2011).

48. Treebak, J. T. et al. Acute exercise and physiological insulin induce distinct phosphorylation signatures on TBC1D1 and TBC1D4 proteins in human skeletal muscle. J. Physiol. (Lond.) 592, 351–375 (2014).

49. Hoffman, N. J. & Elmendorf, J. S. Signaling, cytoskeletal and membrane mechanisms regulating GLUT4 exocytosis. Trends Endocrinol. Metab. 22, 110–116 (2011).

50. Nedachi, T., Fujita, H. & Kanzaki, M. Contractile C2C12 myotube model for studying exercise-inducible responses in skeletal muscle. Am. J. Physiol. Endocrinol. Metab. 295, E1191–1204 (2008).

51. Taylor, E. B. et al. Discovery of TBC1D1 as an insulin-, AICAR-, and contraction-stimulated signaling nexus in mouse skeletal muscle. J. Biol. Chem. 283, 9787–9796 (2008).

52. Vichaiwong, K. et al. Contraction regulates site-specific phosphorylation of TBC1D1 in skeletal muscle. Biochem. J. 431, 311–320 (2010).

53. Cartee, G. D. Of mice and men: filling gaps in the TBC1D1 story. J. Physiol. (Lond.) 588, 4331–4332 (2010).

54. Chung, H. S. et al. Implications of circulating Meteorin-like (Metrnl) level in human subjects with type 2 diabetes. Diabetes Res. Clin. Pract. 136, 100–107 (2018).

55. Lee, J. H. et al. Serum Meteorin-like protein levels decreased in patients newly diagnosed with type 2 diabetes. Diabetes Res. Clin. Pract. 135, 7–10 (2018).

56. Ngoei KRW. et al. Structural Determinants for Small-Molecule Activation of Skeletal Muscle AMPK a2ß21 by the Glucose Importagog SC4. 25, 1–10 (2018).

